# Coding variants identified in diabetic patients alter PICK1 BAR domain function in insulin granule biogenesis

**DOI:** 10.1101/2020.10.05.325951

**Authors:** Rita C. Andersen, Matthew D. Lycas, Jan H. Schmidt, Nikolaj R. Christensen, Viktor K. Lund, Joscha Rombach, Mikkel Stoklund, Donnie S. Stapleton, Gith Noes-Holt, Mark P. Keller, Anna M. Jansen, Rasmus Herlo, Ole Kjærulff, Birgitte Holst, Alan D. Attie, Ulrik Gether, Kenneth L. Madsen

## Abstract

Bin/amphiphysin/Rvs (BAR) domains are positively charged crescent-shaped modules that shape negatively charged curved lipid membranes during membrane remodeling processes. The BAR domain proteins ICA69, PICK1 and arfaptins have recently been demonstrated to coordinate the budding and formation of immature secretory granules (ISGs) at the *trans*-Golgi network. Here, we identify four coding variants in the PICK1 gene from a Danish whole-exome screening of diabetic patients, that all involve change of positively charged residues in the PICK1 BAR domain. All four coding variants failed to rescue the insulin content in INS-1E cells upon KD of endogenous PICK1. Moreover, two variants showed dominant negative properties. Interestingly, *in vitro* assays addressing the BAR domain function suggest that the coding variants compromised membrane binding capacity but increased capacity to cause fission of liposomes.

Live confocal microscopy and super-resolution microscopy further revealed that PICK1 resides transiently on ISGs before egress via vesicular budding events. Interestingly, this egress of PICK1 was accelerated in the coding variants. We propose that PICK1 assists or complements the removal of excess membrane and generic membrane trafficking proteins, and possibly also insulin from ISGs during the maturation process and that the coding variants may cause premature budding possibly explaining their dominant negative function.

## Introduction

Type 2 Diabetes Mellitus (T2DM) is a global health problem affecting more than 400 million individuals (1, 2). T2DM is a complex heterogeneous metabolic disease with an etiology involving a combination of genetic and environmental risk factors. The pathophysiology is characterized by hyperglycemia caused by insulin resistance and impaired insulin secretion, which subsequently lead to β-cell stress and dysfunction (1). It is therefore not surprising, that genes encoding proteins involved in insulin granule biogenesis, such as proprotein convertases and carboxypeptidase E, have been associated with metabolic dysfunction and T2DM (3).

Insulin granules are derived from the regulated secretory pathway (RSP) starting at the trans-Golgi network (TGN) and are formed by packaging proinsulin and other secretory proteins into a nascent bud, followed by a budding process that generates immature secretory granules (ISGs) (4, 5). Subsequent processing, including the cleavage of proinsulin to insulin followed by condensation in response to acidification and Ca^2+^ -influx, leads to the maturation of ISGs to mature secretory granules (MSGs) (6). Moreover, during maturation excess membrane and generic membrane trafficking proteins are removed by budding of vesicles from ISGs, destined for either a constitutive-like secretion, recycling back to the TGN, or to endocytic/lysosomal compartments (7-10). Although the molecular mechanism is unclear, this vesicle budding is thought to be clathrin-dependent, as clathrin and adaptor protein (AP)-1 are detected on ISGs and budding structures while being absent on MSGs (11-14).

The N-Bin/Amphiphysin/Rvs (BAR) domain group of proteins, comprising islet cell autoantigen 69 (ICA69), protein interacting with C kinase 1 (PICK1), and the arfaptins (collectively named IPA BARs) together represent newly discovered modulators of the RSP (15). N-BAR domains constitute a distinct class of dimeric crescent-shaped structures that are flanked by amphipathic helices and associate with high membrane curvature during membrane budding processes via positively charged residues on their concave surface (16-18). Arfaptin 1 stabilizes the nascent budding granules at the TGN by shielding them from the action of ADP-ribosylation factor (ARF)1 in a protein kinase D (PKD)-dependent manner (19), whereas PICK1 as a heterodimer with ICA69 directly assist the budding process from the TGN and regulates maturation of insulin granules (20-22). While the physiological importance of arfaptin’s function remains to be addressed, knockout (KO) studies of PICK1 and ICA69 in mice revealed glucose intolerance as a result of reduced insulin storage and secretion (20-22).

Here, we report coding variants of the IPA N-BAR proteins, identified in a cohort of T2DM patients and focus on four coding variants that cause change of positively charged residues in the PICK1 BAR domain. Functional assessment of the coding variants showed that they failed to rescue the function of PICK1 in insulin granule biogenesis causing reduced insulin storage in INS-1E cells. Two of the coding variants further lead to impaired function of wildtype (WT) PICK1 in a dominant negative manner. The coding variants displayed impaired binding to curved lipid membranes, but surprisingly increased the ability of PICK1 to facilitate liposome fission. Application of 3-dimensional direct STOchastic Reconstruction Microscopy (3D-dSTORM) revealed localization of PICK1 on a subset of insulin granules and further suggested an abscission-like function of PICK1 on insulin granules reminiscent of vesicular budding events. Strikingly, the coding variants accelerate this abscission-like function. We propose that PICK1 egress from ISGs through vesicular budding events during the maturation process assists or complements removal of excess membrane and generic membrane trafficking proteins, and that the coding variants may cause premature budding possibly explaining their dominant negative function.

## Results

### Four missense mutations of positively charged residues in the PICK1 BAR domain were identified in a cohort of Danish T2DM patients

Whole-exome sequencing (WES) was performed on 1000 Danish individuals with a combined phenotype of T2DM, moderate adiposity (BMI >27.5 kg/m^2^) and hypertension (systolic/diastolic blood pressure > 140/90 mmHg or use anti-hypertensive medication) and the patients were age-matched with 1000 healthy control subjects (23). Based on previous indications of the importance of the IPA N-BAR proteins in insulin granule biogenesis, we examined the exomes for coding variants within these proteins (19-22, 24-26). In arfaptin 1, we identified five coding variants in 27 individuals (13 controls/14 patients) and in arfaptin 2 we only identified a single coding variant in two control subjects (Figure 1A). Furthermore, nine coding variants in ICA69 (5 controls/6 patients) and six coding variants in ICA1L (20 controls/15 patients) were identified, most of which had alterations in the unstructured C-termini (figure 1A). Finally, we identified four coding variants in PICK1, all of which were present in patients (1 control/5 patients), and interestingly, all four missense mutations were in the BAR domain and caused an arginine substitution to either glutamine or histidine; Arg158Gln (R158Q), Arg185Gln (R185Q), Arg197Gln (R197Q) and Arg247His (R247H) (Figure 1A). R158Q and R185Q were each identified in one individual with T2DM, R247H was detected in two individuals with T2DM, while R197Q was identified in one individual with T2DM and one control subject (Figure 1A).

**Figure 1.**
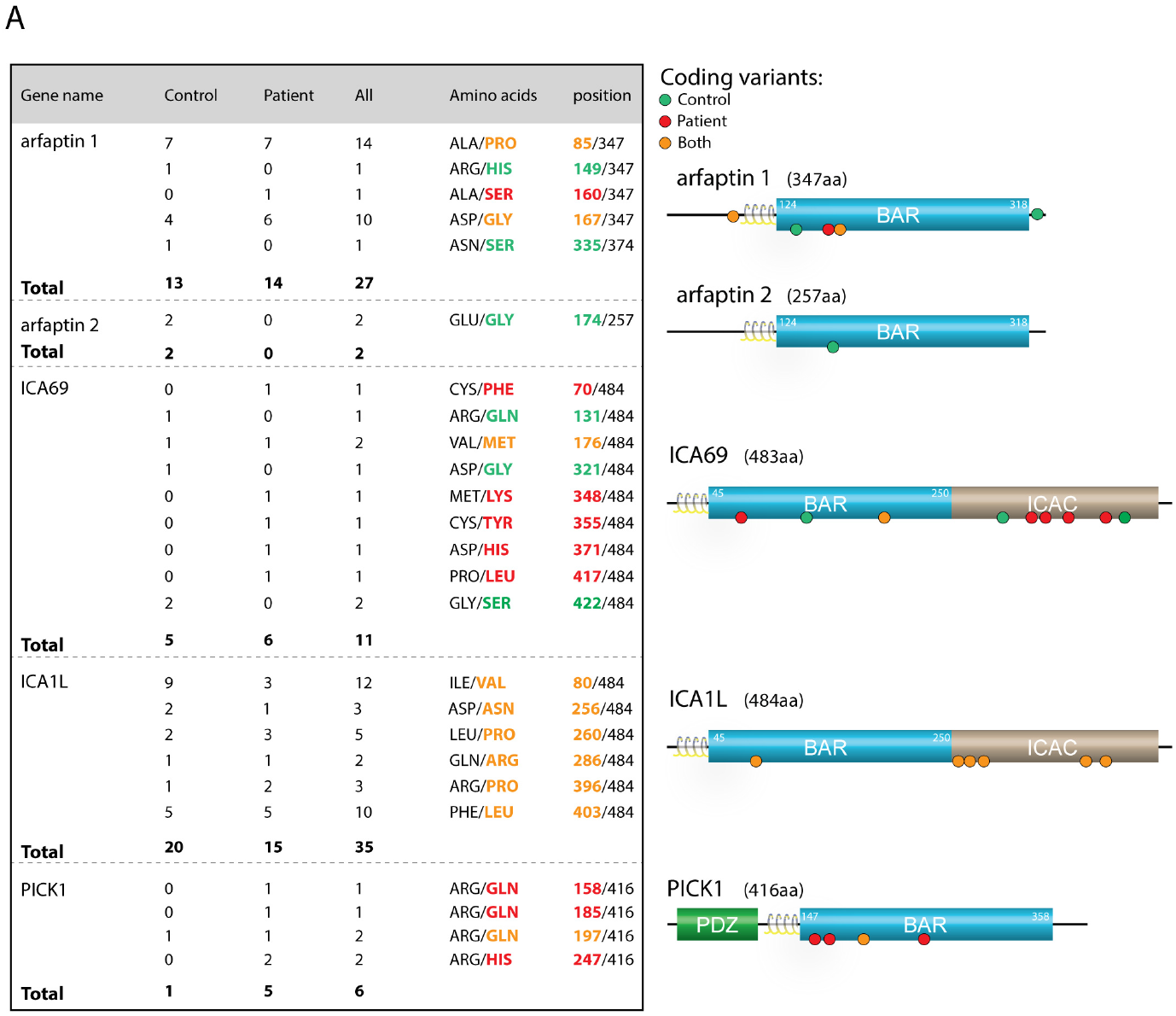
Danish WES of T2DM patients show coding variants in the N-BAR domain proteins. (A) Table of coding variants identified in the N-BAR domain proteins from a Danish WES of T2DM patients and controls. *Right* show the domain organization of the N-BAR domain proteins with position of the coding variants. Green, red and orange dots represent coding variants identified in controls, patients or both, respectively.

Positively charged residues on the concave side of BAR domains have previously been reported to be essential for binding to negatively charged lipids in membranes (16, 27-29). Thus, we hypothesized that the identified coding variants in the PICK1 BAR domain might compromise the membrane binding and deformation capacity of PICK1, causing impaired insulin granule biogenesis in β-cells.

### The coding variants compromise the function of PICK1 leading to reduced insulin content in INS-1E cells

To study the effect of the coding variants in the PICK1 BAR domain on insulin granule biogenesis, we used the insulin-producing INS-1E pancreatic β-cell line. We implemented a molecular replacement strategy as previously described (24, 30), using a lentiviral shRNA construct to knockdown (KD) endogenous PICK1 expression and re-express either eGFP alone (referred to as KD) or eGFP fused to shRNA insensitive PICK1 variants, either WT or the four PICK1 coding variants (referred to as KD + WT, KD + R158Q, KD + R185Q, KD + R197Q, and KD + R247H). A construct expressing eGFP but with deletion of the shRNA was used as a control (referred to as ctrl) (Supplemental Figure 1A). INS-1E cells were transduced with the ctrl, KD, and KD + WT constructs and immunostained for PICK1 and insulin (Figure 2A). Quantification of the PICK1 immunosignal in eGFP positive cells showed a robust decrease in PICK1 expression for KD compared to ctrl, while re-expression of the shRNA insensitive KD + WT construct increased the PICK1 expression level (Figure 2B). Immunoblotting confirmed KD of endogenous PICK1, despite a relatively low transduction efficiency (∼30%), and re-expression of GFP-PICK1 by KD + WT (Supplemental Figure 1, B-C).

**Figure 2.**
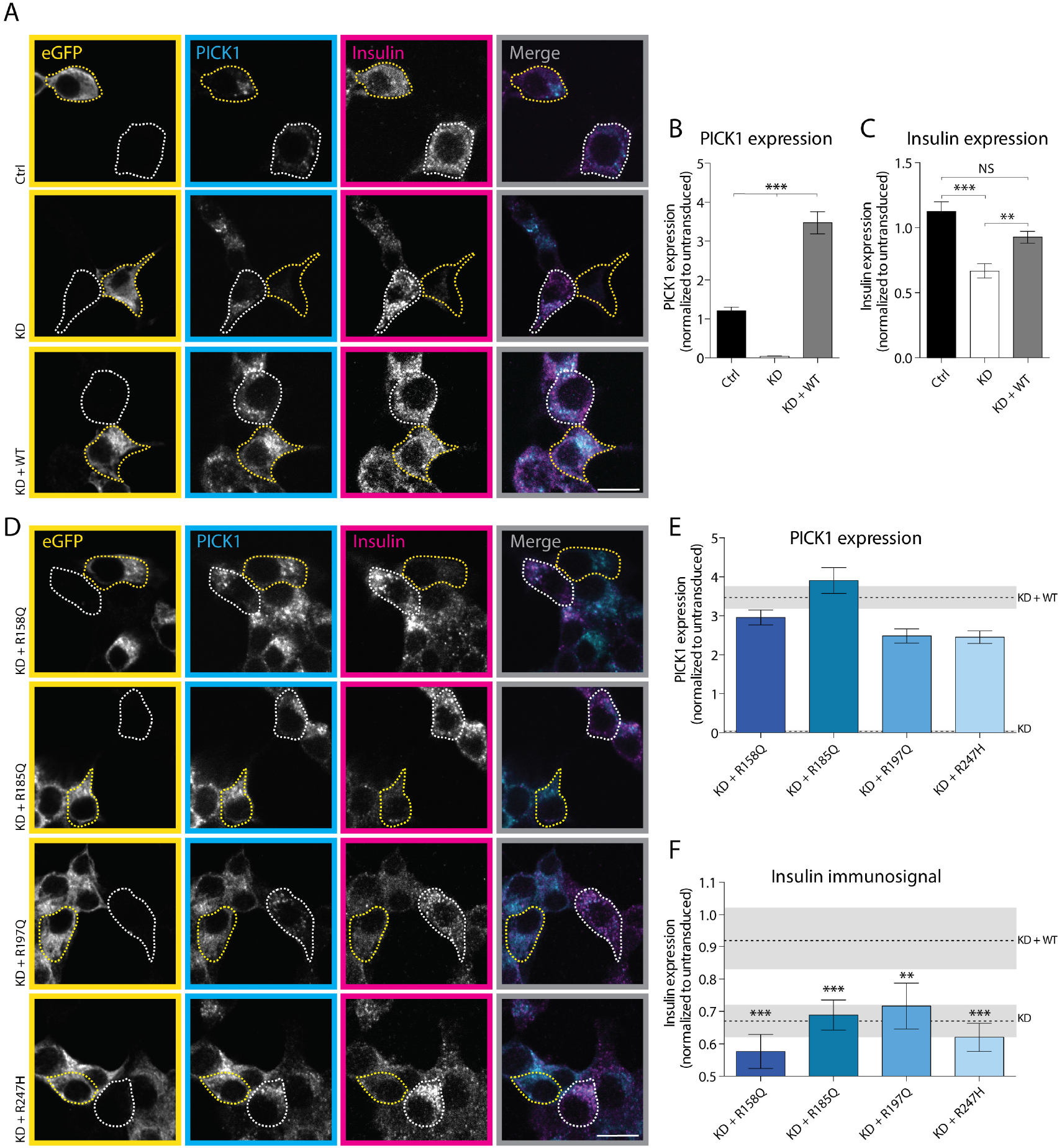
Coding variants in the BAR domain compromise PICK1 function in insulin storage in INS-1E cells. (A and D) Representative confocal images of INS-1E cells transduced with lentiviral constructs as indicated and immunostained for GFP (yellow), PICK1 (cyan) and insulin (magenta). The merged images show the PICK1 and insulin immunosignals. Examples of transduced cells (GFP-positive) are outlined with yellow dotted lines and untransduced cells are outlined with white dotted lines. Scale bar = 10 µm. (A) Representative confocal images of INS-1E cells transduced with ctrl, KD and KD + WT. (B and C) Quantification of the PICK1 and insulin immunosignal from (A). Data are shown as mean ± SEM with Kruskal Wallis test followed by Dunn’s multiple comparisons test, ctrl (n = 122), KD (n = 68) and KD + WT (n = 108). (D) Representative confocal images of INS-1E cells transduced with lentiviral constructs re-expressing PICK1 with each of the four coding variants in the BAR domain. (E and F) Quantification of the PICK1 and insulin immunosignal from (D). The two dashed lines show the mean KD and KD + WT insulin immunosignal ± SEM from (B and C). KD + R158Q (n = 112), KD + R185Q (n = 50), KD + R197Q (n = 64), KD + R247H (n = 102) compared to KD + WT (n = 108), with Kruskal Wallis test followed by Dunn’s multiple comparisons test. Data are shown as mean ± SEM. (\**p*<0.05, \*\**p*<0.01, \*\*\**p*<0.001).

Quantification of the insulin immunosignal in eGFP-positive cells, showed a ∼45% reduction compared to ctrl upon KD of PICK1, whereas KD + WT significantly rescued the insulin expression although not fully to the level of the control (ctrl) (Figure 2C). The same pattern was observed using an enzyme-linked immunosorbent assay (ELISA) (Supplemental Figure 2A), confirming previous studies in both INS-1E cells and isolated islets on the role of PICK1 in insulin granule biogenesis (20, 22, 24).

To examine whether the PICK1 coding variants could rescue the insulin immunosignal upon KD of endogenous PICK1, we quantified the insulin expression from INS-1E cells transduced with the corresponding lentiviral constructs (Figure 2D). Similar to KD + WT (dashed line), the PICK1 immunosignals for the coding variants were increased compared to untransduced cells (indicated by a value of 1) (Figure 2E). Also, localization to the early secretory pathway, as evaluated by colocalization with TGN38, syntaxin 6 and insulin, was intact for the coding variants (Supplemental Figure 3, A-F).

**Figure 3.**
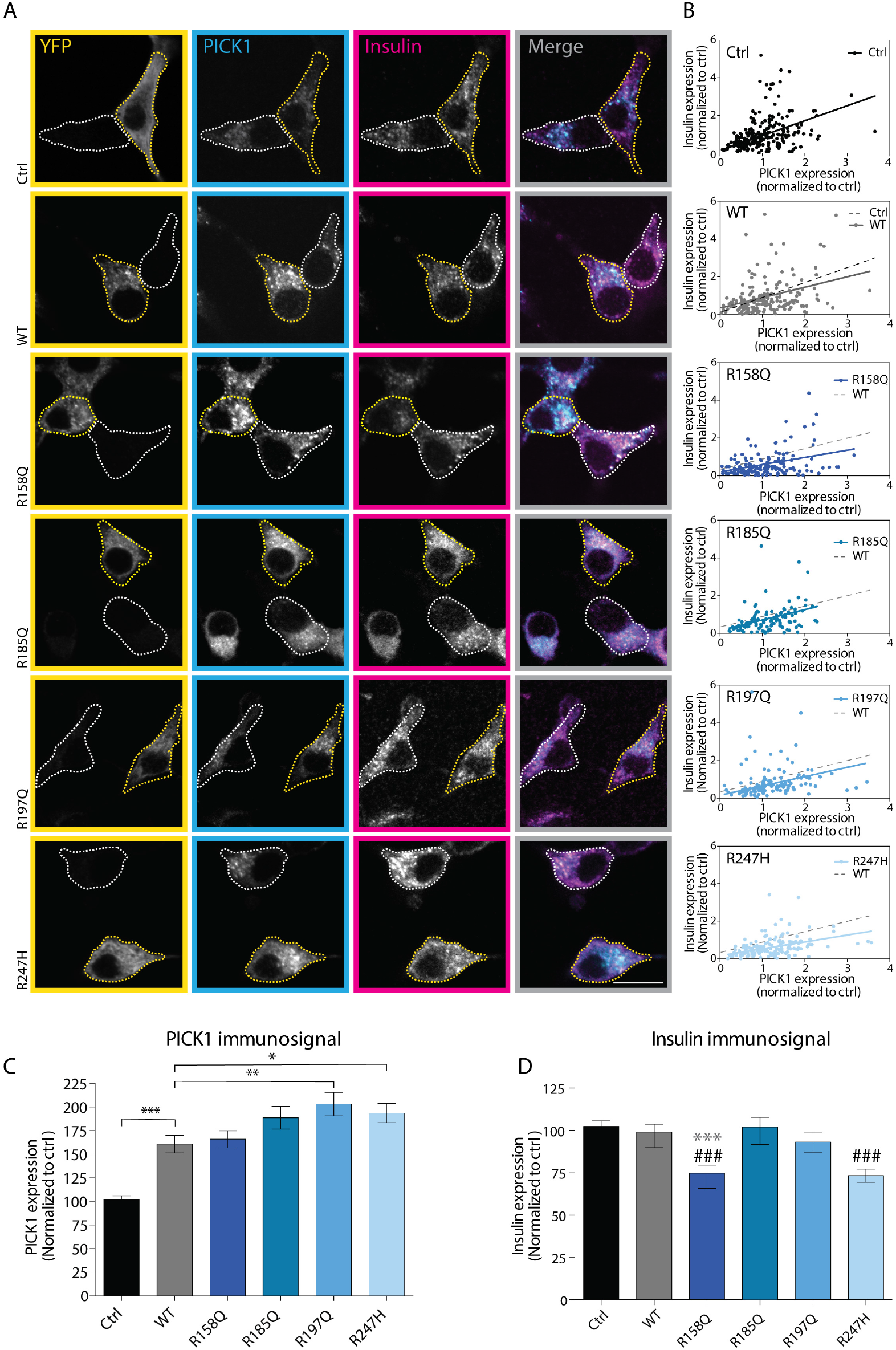
Overexpression of PICK1-R158Q and PICK1-R247H in INS-1E cells reduce insulin storage. INS-1E cells were transiently transfected with YFP alone (ctrl) or YFP fused to PICK1 WT, R158Q, R185Q, R197Q, and R247H. (A) Representative confocal images of INS-1E cells immunostained for YFP (yellow), PICK1 (cyan) and insulin (magenta). Merged images show the PICK1 and insulin immunosignals. Examples of transfected cells (YFP-positive) are outlined with yellow dotted lines and untransfected cells are outlined with white dotted lines. Scale bar = 10 µm. (B) Quantified insulin and PICK1 immunosignals per cell correlate and can be fitted with a linear regression, grey dotted line represent the linear regression from PICK1 WT. Expression of PICK1 R158Q and R247H reduce the slope of the correlation (C) Quantification of the PICK1 immunosignal. R158Q (n = 159), R185Q (n = 119), R197Q (n = 125), R247H (n = 140), ctrl (n= 227) compared to WT (n = 188) with Kruskal Wallis test followed by Dunn’s multiple comparisons test. Data are shown as mean ± SEM. (D) Quantification of the insulin immunosignal. The PICK1 coding variants are compared to WT (*p* value shown with *) and to ctrl (*p* value shown with #) with Kruskal Wallis test followed by Dunn’s multiple comparisons test (\**p*<0.05, \*\**p*<0.01, ***/###*p*<0.001).

Nonetheless, all four PICK1 coding variants failed to rescue insulin expression with significant reductions (∼20-40%) compared to KD + WT (Figure 2F). The failure to rescue the insulin level was confirmed by ELISA, although the effect on insulin levels were slightly less pronounced, since not all cells were transduced (Supplemental Figure 2B).

The decreased insulin content upon KD of PICK1 could be a consequence of unprocessed proinsulin as previously reported (20, 22). However, using ELISA we observed no differences in proinsulin content after KD of PICK1 compared to ctrl or KD + WT (Supplemental Figure 2C), nor did we observe a significant change in proinsulin content after re-expression of the four PICK1 coding variants (Supplemental Figure 2D).

### Overexpression of R158Q and R247H in INS-1E cells decreased the insulin immunosignal in a dominant negative fashion

Since all five patients identified with the coding variants in PICK1 are heterozygous, we next assessed putative dominant negative role of the mutations. To this end, we overexpressed the PICK1 coding variants on top of endogenous PICK1 by transiently transfecting INS-1E cells with YFP-PICK1 WT or either of the four PICK1 coding variants and immunostained for PICK1 and insulin (Figure 3A). As expected, the PICK1 immunosignal in INS-1E cells, overexpressing YFP-PICK1 WT or the four coding variants, was increased compared to ctrl cells (indicated by a value of 100) (Figure 3C). Interestingly, correlations of the quantified insulin immunosignal versus the corresponding PICK1 immunosignal of individual cells suggested a decreased slope for both R158Q and R247H compared to PICK1 WT (Figure 3B), indicating that the two PICK1 coding variants have a negative effect on the insulin content. Quantification of the total insulin immunosignal revealed a significant reduction (by ∼25%) for R158Q and R247H compared to PICK1 WT and/or ctrl (Figure 3D). Neither overexpression of R185Q nor R197Q caused changes in the correlation between insulin and PICK1 or the total insulin immunosignal compared to ctrl or PICK1 WT (Figure 3, B-D). These data suggest that R158Q and R247H not only have reduced functionality, but also can exert a dominant negative effect to suppress the function of WT PICK1.

### The coding variants alter PICK1 clustering and liposome binding

Whereas N-BAR domain proteins in general display a tubular distribution upon overexpression in e.g. COS7 cells, PICK1 (27-30) (similar to e.g. endophilin B1(31)) displays a distinct punctuate pattern upon overexpression. This punctuate distribution has previously been used to assess BAR domain function, and mutations of positive charges in the BAR domain have been shown to compromise this clustering (27-30). Thus, we examined whether our coding variants affected the clustering propensity of the BAR domain using a YFP-Δ1-101PICK1 construct, comprising the PICK1 N-BAR domain region (Supplemental figure 4A). Upon heterologous expression in COS7 cells we observed intense clustering of PICK1 WT as described previously (Figure 4A) (24, 28). Constructs with the coding variants expressed to the same extent as or better than YFP-Δ1-101PICK1 WT (Supplemental figure 4B), but YFP-Δ1-101PICK1 R185Q and YFP-Δ1-101PICK1 R247H were significantly less prone to clustering than YFP-Δ1-101PICK1 WT whereas YFP-Δ1-101PICK1 R158Q and YFP-Δ1-101PICK1 R197Q remained unchanged (Figure 4, A-B). Moreover, R185Q and R247H showed an increase in the fraction of small clusters (<500 nm) at the expense of large clusters (>1000 nm) (Figure 4A).

**Figure 4.**
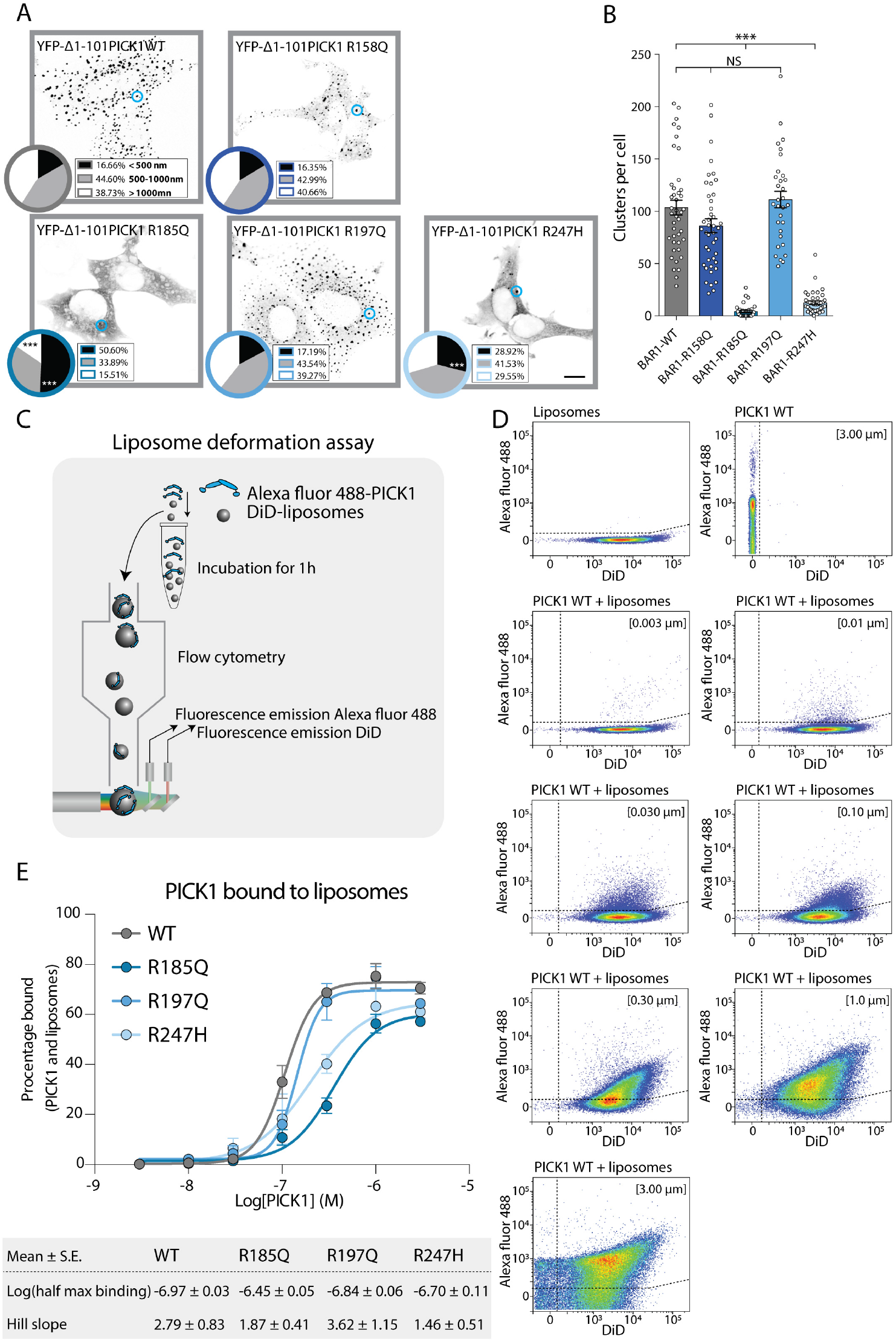
The R185Q and R247H coding variants display impaired liposome binding capacity. (A) COS7 cells were transiently transfected with YFP-Δ1-101 PICK1 WT or the four coding variants. Representative confocal images are shown with an inverted grayscale, where blue circles represent PICK1 clusters. Scale bar = 10 µm. On each image is a diagram showing the fractional size distribution, defined as clusters < 500 nm, 500-1000 nm, and > 1000 nm. R158Q (n = 43), R185Q (n = 32), R197Q (n = 31), R247H (n = 40) compared to WT (n = 41), with two-way ANOVA followed by Dunnett’s multiple comparisons test. (B) The quantification of clusters per cell. Kruskal Wallis test followed by Dunn’s multiple comparisons test. (C) Experimental design of single liposome binding of protein to curved membranes by flow cytometry. (D) Representative two-parameter density plot of primary data output from the liposome binding assay after 1 h incubation showing fluorescent intensities (A.U.) of AF488 (PICK1) vs. DiD (liposomes) for samples containing liposomes or PICK1 alone as well as liposomes with increasing concentration of PICK1 WT (3 to 3000nM). Densities of observations indicated from low (blue) to high (red). (E) Dose-response curves for total binding of PICK1 WT and the PICK1 coding variants to liposomes, shown as the fraction of total events above the two gates indicated by dashed lines (see Methods). The half max binding and hill slope values were determined by nonlinear regression fits. Data are shown as mean ± SE, n = 3 individual experiments.

To directly assess the effect of the coding variants on the association of PICK1 with curved membranes, we employed a modified version of the previously described Single Liposome Curvature Sensing (SLiC) assay (32) adapted to a high-throughput analysis by flow cytometry (Figure 4C).

To detect binding of PICK1 to liposomes, we applied increasing concentrations of purified PICK1 labeled with Alexa Fluor 488 (3 to 3000nM) to a constant concentration of small unilamellar liposomes (SUVs), prepared from bovine brain extract (Folch fraction 1, ∼2.5 µg/mL) and labeled with the lipophilic dye (DiD) followed by incubation for 1 h (Figure 4C). We determined the dual fluorescence intensities by flow cytometry and defined binding, as events passing two gates derived from intensities observed in samples with liposomes and PICK1 WT alone (Figure 4D, top panel *left* and *right*, respectively). The fraction of protein-bound liposomes accumulated with increasing concentration of PICK1 WT giving rise to a sigmoidal binding curve with a half max binding of ∼100 nM (−6.97 ± 0.03) and a hill coefficient of 2.8 indicative of cooperative binding (Figure 4E), similar to our previous findings for endophilin and amphiphysin in the traditional SLiC assay (18). PICK1 R185Q, R197Q, and R247H purified equally well to PICK1 WT (Supplemental Figure 4C), and the PDZ domain function was unaltered testing the overall integrity of the protein (Supplemental Figure 4, D-E). Unfortunately, we were unable to purify PICK1 R158Q. We similarly performed concentration-dependent flow cytometry experiments for the PICK1 coding variants (Supplemental Figure 5, A-C). Quantification of the binding curves showed that R197Q displayed a binding curve similar to PICK1 WT, whereas both R185Q and R247H displayed lower hill coefficients as well as higher concentrations to reach half maximal binding, indicating weaker and less cooperative binding (Figure 4E).

**Figure 5.**
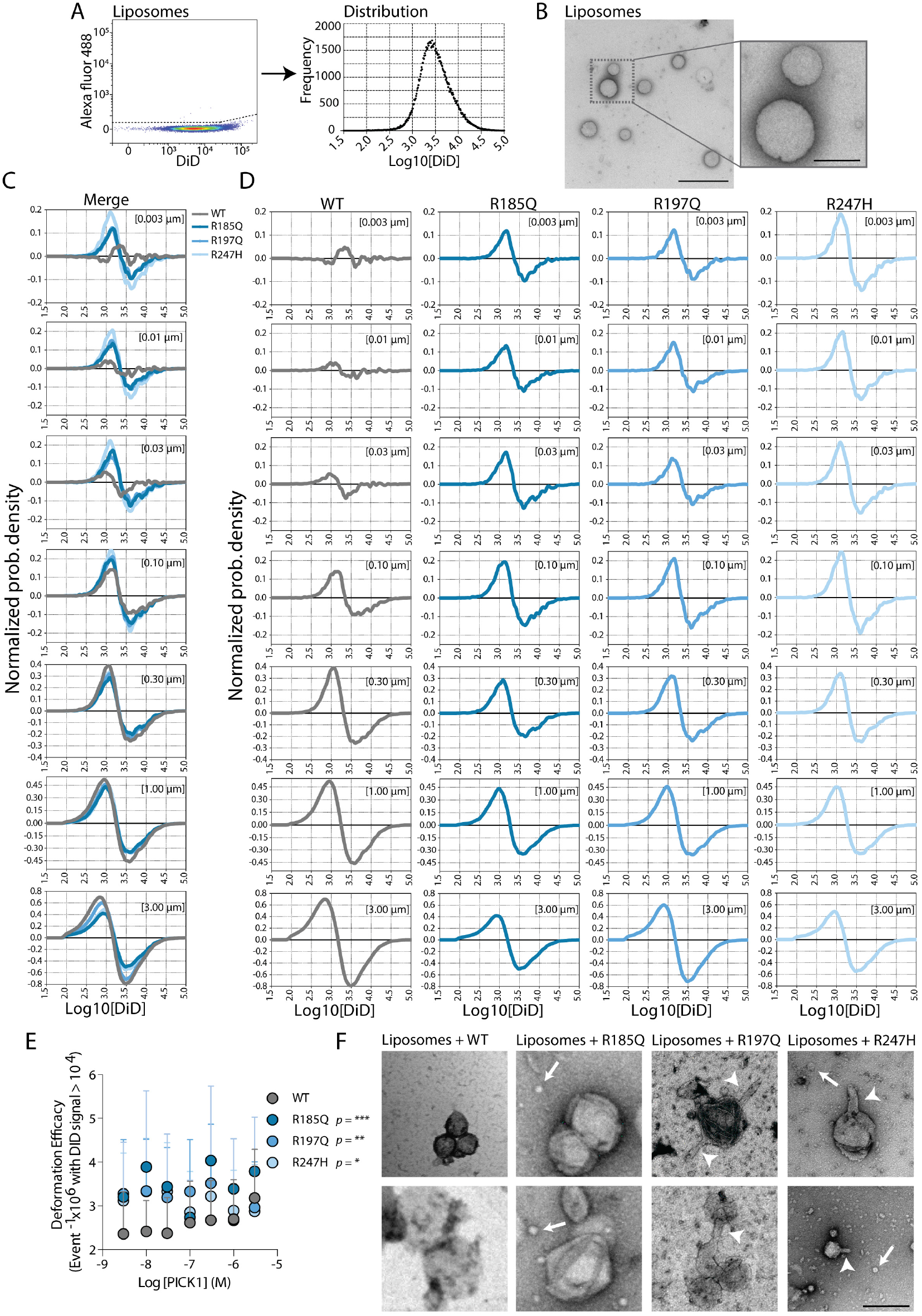
The PICK1 coding variants show increased fission efficacy at lower concentration and altered liposome deformation. (A) Frequency distribution of the DiD intensities extracted from figure 4D indicating the size distribution of DiD labelled liposomes prior to incubation with protein. (B) Representative negative-stained EM image of liposomes, scale bar = 1 µm. *Right* shows insert with higher magnification, Scale bar = 200 nm. (C) Representative flow cytometry experiment showing change in frequency distribution of liposomes (normalized kernel density estimations of DiD fluorescence compared to control) upon incubation (1 h) for a concentration range (0.003 to 3 uM) of PICK1 WT and the PICK1 coding variants as a measure of liposome fission efficacy. (C) Same curves as in (B) shown individually for the constructs. (D) Liposome fission efficacy quantified as loss of event with DiD signal > 10^4^, n = 5 individual experiments. (E) Representative negative-stained EM images of liposomes incubated with purified PICK1 WT, R185Q, R197Q and R247H, respectively. Arrowheads point to tubular structures while arrows point to small liposome structures. Scale bar = 200 nm.

We have recently demonstrated that membrane-curvature dependent binding of PICK1 WT to SUVs is dependent on the amphipathic helix preceding the BAR domain as this binding is strongly reduced by mutational disruption of the helix (24). The coding variants identified in this paper, however, did not display impaired membrane curvature sensing (Supplemental Figure 5, D-E).

### The coding variants potentiate the liposome fission efficacy of PICK1

Recent studies have demonstrated potent fission capacities of BAR domain proteins, challenging the current consensus of BAR domain proteins as general stabilizers of tubular membrane structures (33, 34). Indeed, from our primary data of the protein-liposome interaction by flow cytometry, it was evident that the liposome fluorescence was gradually shifted towards lower intensities with increasing concentration of PICK1, indicative of liposome fission (Figure 4D). To visualize this change in distribution, we used kernel density estimates of the fluorescent intensities of liposomes. Normalization of density estimates after incubation with increasing concentration of PICK1 to the initial liposome distribution (Figure 5A and B) revealed an increase in frequency of liposomes with Log DiD intensities from 2.5-3.3 at the expense of liposomes with Log DiDintensities > 3.3. (gray, Figure 5, C and D). This shift in the size distribution, which represent liposomes fission, was accentuated with increasing concentration of PICK1 (gray, Figure 5, C and D). Surprisingly, the coding variants displayed more potent liposome fission at non-saturation concentrations of proteins (<1 µM) and consequently less concentration dependence (Figure 5, C-E). The overall shapes of the normalized density changes were identical for PICK1 WT and the coding variants (Figure 5,C) implying that only the efficacy but not the size dependence of fission was affected by the coding variants.

To directly visualize the fission products resulting from incubation of liposomes with purified PICK1, we turned to transmission electron microscopy (TEM). The images revealed numerous adherent, small to medium-sized liposomes (50-200 nm) (Figure 5F) that were not observed in absence of PICK1 (Figure 5B). This suggests that PICK1 is prone to perform abscission of larger liposomes into small to medium-sized liposomes rather than tubulation as seen for most N-BAR domains (16). R185Q and R197Q likewise formed adherent liposome structures, but R185Q also produced numerous, small liposome structures, as did PICK1 R247H. Interestingly, R197Q, and in particularly R247H, also formed tubular membrane structures akin to the structures observed with other N-BAR domains. Notably, we never observed tubular structures for PICK1 WT or R185Q.

Inspection of our all-atom molecular dynamics simulation model based on the homologous arfaptin 2 BAR domain (35), suggests that R158, R185, R197 and R247 are all engaged with acidic residues within the BAR domain, likely serving to stabilize helix-helix interactions within the coiled-coil structure (Supplemental Figure 6A). This suggests that the functional effect on membrane binding and deformation may rather result from structural reorganization or changes in oligomerization propensity on liposomes rather than a simple loss of positive charge.

**Figure 6.**
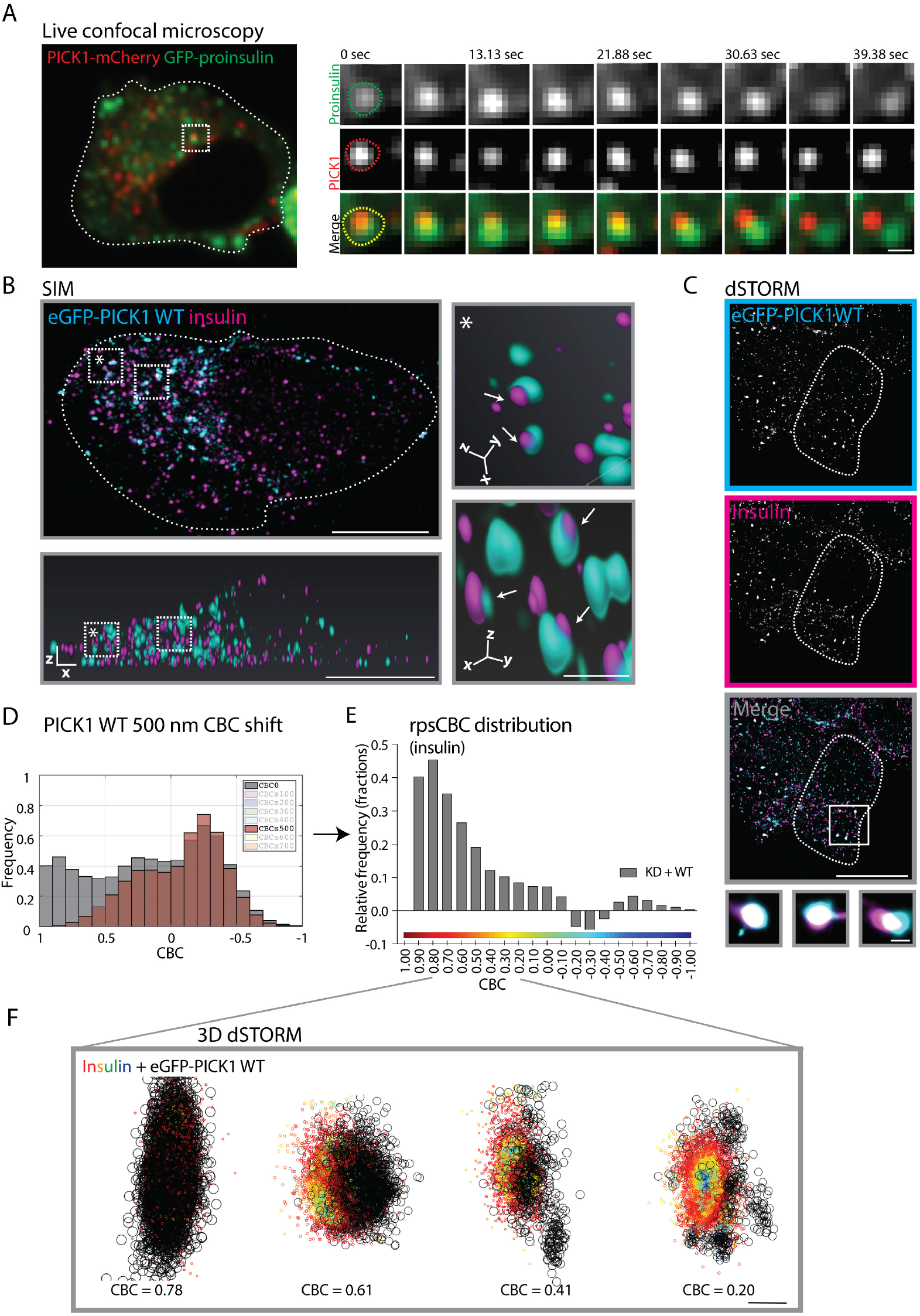
PICK1 resides transiently on insulin ISGs before budding off. (A) GRINCH cells were transiently transfected with PICK1-mCherry. Representative GRINCH cell shows a coocalized GFP-proinsulin (green) and PICK1-mCherry (red) cluster indicated by the square. *Right* shows the time-lapse imaging of merged z-stack (500 nm) acquired over time during steady-state condition (11 mM glucose). Upper two rows present a PICK1-mCherry positive cluster and a GFP-proinsulin cluster, both shown in grey scale. The third row shows merge images. Time is in seconds and scale bar = 1 µm. (B) Representative SIM image of INS-1E cells transduced with KD + WT, and immunostained for PICK1 (cyan) and insulin (magenta). *Bottom* same INS-1E cell in 3D re-construction, scale bar = 5 µm. *Right* shows insert with higher magnification of overlapping PICK1 and insulin granules (arrow). Scale bar = 500 nm. (C) Representative dSTORM image of INS-1E cells transduced with KD + WT, and immunostained for eGFP-PICK1 (cyan) and insulin (magenta), scale bar = 5 µm. *Bottom* shows the insert with higher magnification of overlapping PICK1 and insulin granules. Scale bar = 250 nm. (D) Insulin CBC shift analysis. The PICK1 clusters were shifted +500 nm in the x direction, and the CBC distribution of the insulin granules are recalculated (brown) and overlaid on the original CBC distribution (grey). (E) Shows the difference in CBC between the original from (B) (grey) and the +500 nm shifted (brown). this we refer to as random proximity subtracted CBC (rpsCBC) distribution. Note that many points are not assigned a CBC values (NA) when shifted. n = 5 individual experiments. (F) The 3D images display distinctive colocalized clusters of insulin (colored by CBC scale) and eGFP-PICK1 WT (black), ordered by CBC values ranging from 0.78 to 0.20 and indicative of PICK1 fission from insulin granules. Scale bar = 200 nm.

### PICK1 resides transiently on insulin ISGs before budding off during maturation

Next, we assessed whether PICK1-dependent abscission of liposomes might relate to its function in dense core vesicle biogenesis. To evaluate the dynamic association of PICK1 with secretory granules in living cells, we took advantage of the glucose-responsive, insulin-secreting, C-peptide-modified human proinsulin (GRINCH) INS1 cell line that stably expresses eGFP-proinsulin (36). Live confocal microscopy of GRINCH cells transiently transfected with PICK1-mCherry demonstrated a partial overlap of the signal from PICK1-mCherry and eGFP-proinsulin (Figure 6A and Supplemental Figure 7A), which is in good agreement with our immunostainings (Supplemental Figure 3, A-B) and previous live microscopy studies with PICK1 and phogrin (20). Interestingly, by following the dynamics of individual puncta of PICK1-mCherry and eGFP-proinsulin clusters in GRINCH cells, we often observed separation of the colors over time indicating that the PICK1 association with insulin granules was of extended, but ultimately transient nature (Figure 6A, Supplemental Figure 7 B-C and Video 1).

**Figure 7.**
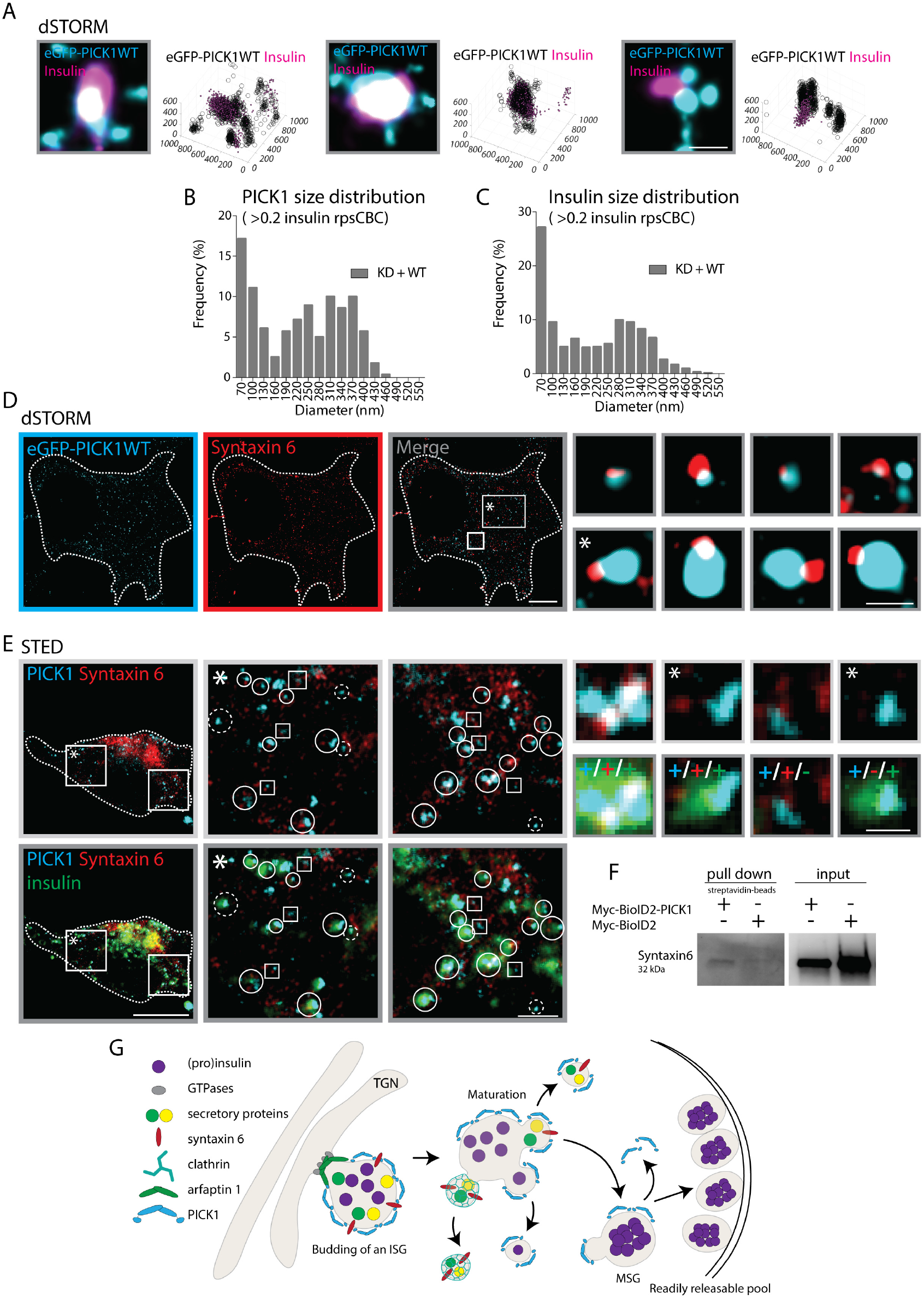
Super resolution imaging implicates PICK1 in novel egress route from ISGs. (A) Representative dSTORM images of transduced INS-1E cells. Several examples demonstrate PICK1 (cyan) in small clusters around insulin granules (magenta) in 2D. Scale bar = 250 nm. Same data illustrated in 3D displaying small amounts of insulin (magenta) in the small surrounding structurs of PICK1 (black). Axis indicate nm. (B-C) The size distribution of PICK1 and insulin clusters confirms high prevalence of small clusters < 100nm in the colocalized structures defined as CBC > 0.2 insulin from the rpsCBC distribution. (D) Representative dSTORM image of INS-1E cells transduced with KD + WT, and immunostained for eGFP-PICK1 (cyan) and syntaxin 6 (red), scale bar = 5 µm. *Right* shows insert with higher magnification of small overlapping PICK1 and syntaxin 6 positive structures. Scale bar = 250 nm. (E) Representative STED image of INS-1E cells immunostained for PICK1 (cyan), syntaxin 6 (red) and insulin (green), scale bar = 5 µm. *Top* and *bottom* panels show the inserts with higher magnification with PICK1/syntaxin 6 and PICK1/syntaxin 6/insulin immunostaining, respectively. Squares represent colocalized PICK1/syntaxin 6 clusters, dashed circles represent colocalized PICK1/insulin clusters and whole circles represent colocalization between PICK1/syntaxin 6/insulin clusters, scale bar = 1 µm. *Right* is higher magnification of the colocalizations, scale bar = 250 nm (F) INS-1E cells were transient transfected with a Myc-BioID2-PICK1 construct or Myc-BioID2 as control. Biotinylated proteins were pulled down from cell lysates with streptavidin-beads for immunoblotting against syntaxin 6. n = 3 individual experiments. (G) Putative model for the role of the IPA N-BAR domain proteins in insulin granule biogenesis. Arfaptin 1 controls the neck of growing ISGs at the TGN in a complex with effector and kinases, while PICK1, either in a homo- or heterodimeric complex with ICA69, localizes around the growing ISGs promoting membrane fission. After the ISGs are budded off, the ISGs undergo a maturation process including the cleavage of proinsulin into insulin followed by condensation. We propose that PICK1, during multiple budding events remove excess membrane and cargo from the insulin granules in a process complimentary to the clathrin dependent egress.

Interpretation of live microscopy can be obscured by rotation and overlay in the z-axis within the confocal slices, so to further examine the association between PICK1 and insulin granules, we turned to 3D-resolved structured illumination microscopy (SIM). We transduced INS-1E cells with KD + WT and immunostained for eGFP(-PICK1) and insulin, and assessed the spatial distribution (Figure 6B). Again, we observed overlapping structures between PICK1 and insulin throughout the cell, although most prominently in the perinuclear region. Moreover, when zooming in on individual granules numerous structures were observed with either partial overlap of the signal or side-by-side localization consistent with transient structures in a fission process (Figure 6B).

### 3D-dSTORM enables quantification of PICK1 budding from insulin granules

To increase resolution and enable better visualization as well as quantification of the localization of PICK1 in relation to insulin granules and thereby the putative budding process, we turned to dual color 3-dimensional direct STOchastic Reconstruction Microscopy (3D-dSTORM). We transduced the INS-1E cells with KD + WT and immunostained for eGFP(-PICK1) and insulin (Figure 6C). We used the insulin signal to define the size of the insulin granules and the eGFP signal to identify PICK1 positive clusters (described in Methods and (24)). We observed many examples of full overlap between the PICK1 and the insulin localizations as described previously, however, we also observed insulin clusters with all PICK1 localizations skewed to one side as well as insulin clusters depleted in PICK1 localizations, but with one or multiple adjacent clusters of PICK1 localizations down to 50 nm in size (Figure 6, C,F and 7A).

To quantitatively describe the degree of overlap between insulin granules and PICK1 clusters we next took advantage of coordinate-based colocalization (CBC) analysis with a value of 1 indicating a perfect overlap while −1 represents no overlap. To address which range of CBC values reflected a biologically relevant overlap as opposed to random proximity, we shifted the two channels with respect to each other in steps of 100nm (from 100-700 nm), and subtracted the resulting CBC histograms to derive random proximity subtracted CBC (rpsCBC) histograms (Figure 6, D-E and Supplemental Figure 8, rpsCBC histograms in grey). Shifts beyond 500 nm did not change the CBC histograms further, and consequently all rpsCBC histograms are derived as original CBC histogram subtracted by the 500 nm shifted histograms. Importantly, high CBC values (> 0.5) are significantly enriched in the rpsCBC relative to the histogram with all CBC values, indicating that these values are non-random, whereas values below 0 are depleted in accordance with their predicted random nature. The full rpsCBC histogram for PICK1 WT range continuously from 0.9, indicating almost full overlap, down to 0, indicating weak proximity, with a minor local maximum at 0.8 (Figure 6E). To visualize exemplary structures corresponding to different CBC values, representative 3D reconstructed dSTORM images of colocalized PICK1 (black) and insulin (rainbow gradient) clusters are shown with the insulin CBC values ranging from CBC = 0.78 reflecting a complete overlap, over CBC = 0.61, which reflect a one-sided assembly of PICK1, to CBC = 0.41 reflecting PICK1 being partially dissociated from the insulin granule and finally to CBC = 0.20, which reflect the smaller PICK1 clusters surrounding the insulin granule (Figure 6F). Tentative arrangement of such combined insulin/PICK1 structures according to the CBC values convey the impression of structures that are coated by PICK1, and which buds off insulin granules, which is in accordance with our live confocal microscopy (Figure 6A, Supplemental Figure 7, and video 1) and SIM images (Figure 6B).

**Figure 8.**
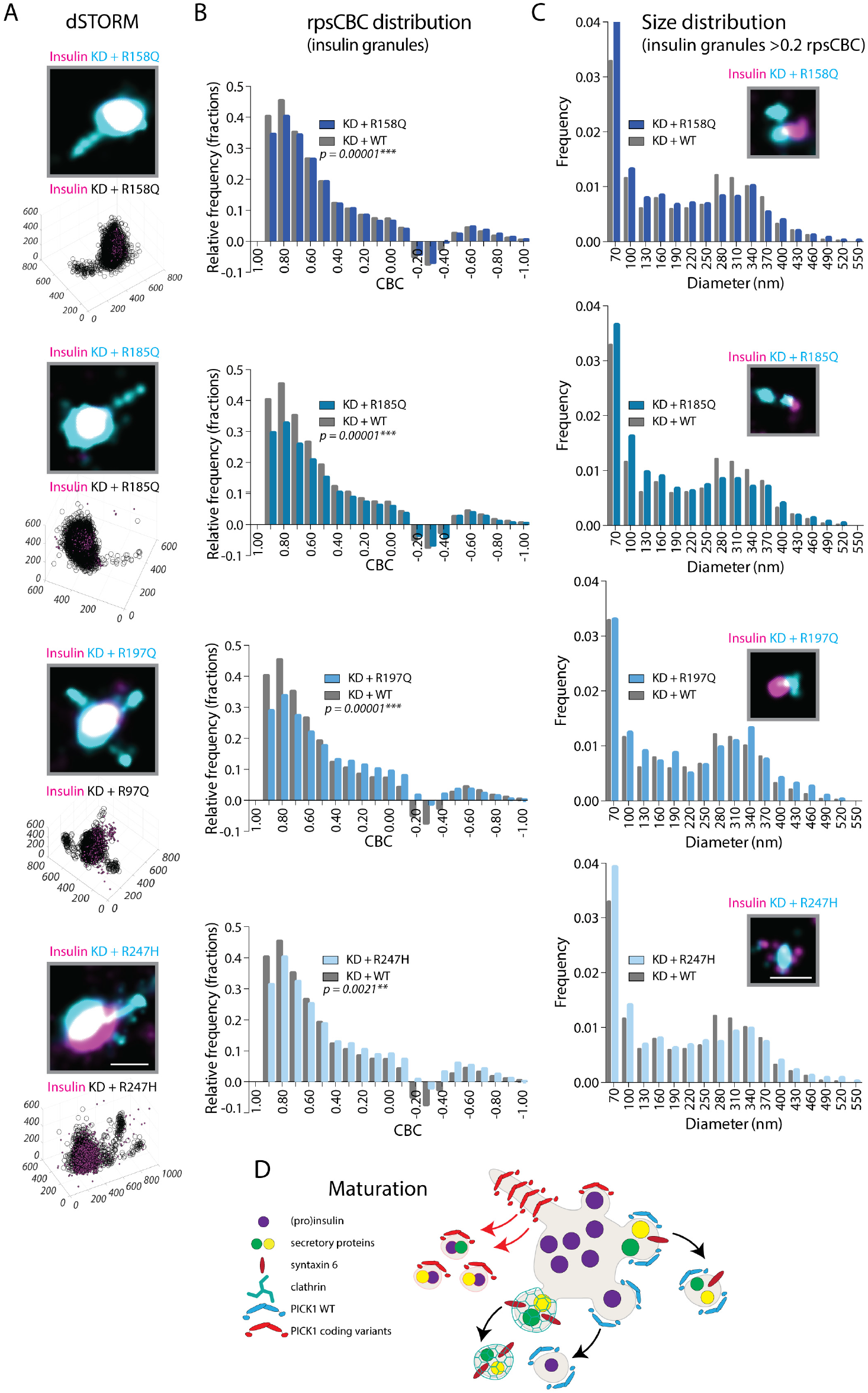
The PICK1 coding variants alter fission from insulin granules. INS-1E cells were transduced with the lentiviral constructs KD + WT and coding variants. (A) Representative 3D dSTORM images of PICK1 (black) in tubular structures colocalized with insulin (magenta). Scale bar = 250 nm. (B) The rpsCBC distribution of PICK1 clusters to insulin granules of the four coding variants (shades of blue) compared to PICK1 WT (grey). Kolmogorov-Smirnov test was used to test the cumulative distribution, n = 4-5 individual experiments. (\*\*\**p*<0.001). (C) The size distribution of colocalized insulin granules and PICK1 clusters, defined as rpsCBC > 0.2, shown with representative dSTORM images of small insulin clusters (magenta) colocalized with PICK1 (cyan). Scale bar = 250 nm. (D) Proposed model for the PICK1 coding variants in insulin granule biogenesis. We propose that the PICK1 coding variants, with increased abscission efficacy, may cause tabulation and premature budding from the SGs during and/or after the maturation process generating small clusters that contain excess membrane cargo and insulin.

### Super resolution imaging implicates PICK1 in novel egress route from ISGs

We next looked into the PICK1 clusters adjacent to insulin granules, which were mostly less than 100 nm in diameter. In many cases, the visual inspection strongly resembled multiple budding processes originating from the same ISG and in several cases these structures also showed low number of insulin localizations (Figure 7A). To obtain support that these structures were not randomly distributed in proximity to ISG, we extracted the size distribution of PICK1 (Figure 7B) and in insulin clusters (Figure 7C) from the part of the rpsCBC histogram (Figure 6E) with values > 0.2, which amounted to 9.2% of all the insulin granules (1191 of 12994 total) for PICK1 WT. Indeed, both PICK1 and insulin structures displayed a bimodal distribution with a significant fraction of small structures (>100 nm) consistent with a budding process involving PICK1 and insulin.

We next considered the possibility that these PICK1 positive buds on the insulin granules may represent precursors of vesicles responsible for the removal of excess membrane and generic membrane trafficking proteins, such as syntaxin 6, during the maturation process (13, 37). We transduced INS-1E cells with our lentiviral construct enabling molecular replacement of endogenous PICK1 with eGFP-PICK1 (KD + WT) and immunostained for eGFP(-PICK1) and syntaxin 6, and again used dSTORM to evaluate the putative overlay. Indeed, we observed colocalization of small (<100 nm) PICK1 and syntaxin 6 clusters (Figure 7D, top zooms and Supplemental figure 9A), consistent with the two proteins budding of ISGs together. However, we also frequently observed small syntaxin 6 clusters on one side of larger PICK1 clusters (∼150-200 nm) (Figure 7D, bottom zooms and Supplemental Figure 9B), resembling syntaxin 6 positive budding processes devoid of PICK1. Consistently, we also observed small (<100nm) syntaxin 6 clusters without PICK1 (not shown).

To verify that these structures were indeed associated with insulin granules, we turn to STimulated Emission deletion (STED) microscopy. This enabled us to perform two-color super resolution (PICK1 and syntaxtin 6) together with confocal resolution (stained for insulin). We indeed observed numerous insulin granules positive for both PICK1 and syntaxin 6 (Figure 7E, white circles), and in some cases PICK1 and syntaxin 6 signal was overlapping (zoom (i)) and in other clearly separated (zoom (ii)). We also observed several insulin granules with PICK1 but no detectable syntaxin 6 (Figure 7E, white dashed circles). Finally, SIM revealed clathrin association with a subset of PICK1 positive structures, but only with partial overlap (Supplementary Figure 9C).

To biochemically probe the proximity of PICK1 and syntaxin 6, we took advantage of the proximity-dependent biotin identification (Bio-ID2) approach, which uses BirA to biotinylate proteins in close proximity to a bait protein (38, 39). PICK1 was fused with the Myc-BioID2 construct, as bait, and INS-1E cells were transiently transfected with the Myc-BioID2-PICK1 construct. Immunoblotting confirmed the presence of syntaxin 6 in streptavidin pull-down (Figure 7F), suggesting that PICK1 and syntaxin 6 are indeed in close spatial proximity in INS-1E cells. Similar blotting for clathrin did not show difference in pull-down between cells transfected with Myc-BioID2 and Myc-BioID2-PICK1 (Supplementary Figure 9D).

In summary, we propose that PICK1 serves a role in egress of vesicles from ISGs carrying small amounts of insulin and generic membrane trafficking proteins. This process appears complementary to the clathrin dependent egress, which is responsible for removing excess membrane and generic membrane trafficking proteins during the maturation process of ISGs (Figure 7G).

### The coding variants in the PICK1 BAR domain increase fission from insulin granules

Next, we expressed each of the coding variants fused to eGFP and inspected 3D dSTORM images of the transduced INS-1E cells. Similar to PICK1 WT, the coding variant localized to insulin granules (consistent with confocal data Supplementary Figure 3 A and B) and also showed PICK1 clusters adjacent to and surrounding the insulin clusters (Figure 8A). Notably, the coding variants were more prone to produce tubular extensions from the granules, than was the case for PICK1 WT (Supplemental Figure 10, A-E), and these structures occasionally also contained insulin (Figure 8A, Supplemental Figure 10, B-E).

We quantitatively assessed whether the coding variants changed the association between PICK1 and insulin granules, by comparing the rpsCBC histograms for insulin granules in INS-1E cells expressing the PICK1 coding variants to PICK1 WT. Interestingly, the rpsCBC histograms (Figure 8B, shades of blue) for all four coding variants differed significantly from the PICK1 WT rpsCBC histogram (Figure 8B, grey) and showed a reduction in prevalence of insulin granules with the highest CBC values (0.7-0.9) in comparison to PICK1 WT (Figure 8B). In accordance, although less pronounced, we observed an increase in CBC values ranging from 0 to ∼0.4 for R197Q and R247H. These data suggest a change in the dynamic association of PICK1 with insulin granules, with the steady state shifted towards lower overlap, which in turn might reflect increased fission or abscission of vesicles from insulin granules.

To further address whether the PICK1 coding variants might affect fission of insulin granules, we examined the size distribution of the colocalized insulin granules at CBC ≥ 0.2 (Figure 8C). Indeed, we observed an increase in the smallest (≤100nm) PICK1-associated insulin structures for the R158Q, R185Q and R247H coding variants compared to PICK1 WT. These results are in agreement with our *in vitro* studies and indicate that the PICK1 coding variants, surprisingly, might increase the rate of vesicle budding from the insulin granules, consequently generating more small insulin positive clusters (Figure 10, B-E).

Finally, to address how general this phenotype might be, we mimicked the most prominent of the coding variants (R247H) in the *drosophila* PICK1 (dPICK1 K249H-HA). Immuno-labeling of dPICK1 WT-HA in large peptidergic cells in the ventral nerve cord of pupal flies showed localization overlapping with and bordering a GFP Golgin245 suggesting localization at and proximal to the Golgi compartment (Supplemental Figure 11, *top*). Strikingly, the dPICK1 K249H-HA construct clearly dissociated from the GFP-Golgin245 in bright spots that in some case shower distinct tubular shapes (Supplemental Figure 11, *bottom*), suggesting that the R247H coding variant interfere with a structural and functional feature of the PICK1 BAR domain that is preserved across species and may affect the early RSP in many (neuro)endocrine cell types.

## Discussion

The IPA group of N-BAR domain proteins have recently been implicated in insulin granule biogenesis, presumably regulating and inducing membrane deformation of SGs both during the fission of nascent TGN membrane, leading to generation of ISGs, and during the maturation process leading to formation of MSGs (19-22). Here, we report four coding variants in the PICK1 BAR domain that were identified in a WES of Danish T2DM patients. All four missense mutations display compromised function in relation to SG biogenesis in insulin-producing cells with two of the mutations also showing a functional dominant negative effect on the WT protein. In line with the membrane binding properties of the BAR domain, the coding variants caused subtle changes to the interaction of PICK1 with liposomes. For two of the variants, liposome binding strength and cooperativity were reduced, but surprisingly membrane fission/abscission efficacy was *increased*. This prompted us to probe the dynamics of PICK1 association with ISGs by use of both live confocal microscopy and super-resolution microscopy. Our data demonstrated that association with insulin granules is transient consistent with PICK1 showing a steady state association with ∼10% of insulin granules. Further our data supported a role of PICK1 in an egress process from ISGs through vesicular budding events that assists or complements clathrin dependent removal of excess membrane and generic membrane trafficking proteins. Finally, quantitative colocalization analysis of super-resolution data, suggested that the steady-state distribution of the PICK1 coding variants shifted away from full coverage of insulin granules and toward budding structures and small granules consistent with an increased fission/abscission efficacy.

It is clear that PICK1 dysfunction is not a major determinant in the development of T2DM. However, although only 5 of the 1000 T2DM patients in the study had coding variants in the PICK1 gene, disruptive mutations may in rare cases contribute to the disease. The within-study odds-ratio was 5, but it must be emphasized that the power of the study was too low (p = 0.22, Fisher’s exact test) to conclude a significant association between T2DM and BAR domain coding variants, even if they phenotypically are considered as one. The same is true for the two dominant negative variants, R158Q and R247H, considered together (p = 0.12, Fisher’s exact test), however, both of these variants are rare SNPs with a frequency of A=0.00003 (GnomAD_exome and TOPMED) for R158Q and A=0.00002 (GnomAD_exome and TOPMED) for R247H, suggesting that the likelihood for random hits in our sample of 1000 patient is very low. Taken together, higher-powered studies are needed to determine, whether coding variants in the PICK1 BAR domain are indeed part of the complex genetic makeup predisposing to T2DM or possible other PICK1 related pathological conditions.

The causal relation to disease aside, the compromised function of the coding variants in insulin storage provided new and potentially important insight into the molecular function of BAR domain proteins and in particular the functional role of PICK1. Relatively few disease-related mutations in BAR domains have been characterized previously see e.g. (40, 41) and consequently, the current understanding of BAR domain structure function reflects hypothesis-driven mutations of positively charged residues, which show reduced liposome binding and altered cellular localization (16, 27, 42). Two of the mutated arginines (R185Q + R247H) in our study compromised localization in COS7 cells, however, their overall localization to the RSP in INS1E cells was intact. The same two mutations likewise reduced liposome binding, which was associated with a reduced hill coefficient and an increase in fission/abscission efficacy. This is rather surprising given that increased binding and deformation are usually considered interdependent (43-45). Interestingly, our TEM studies demonstrated that membrane deformation by PICK1 WT primarily resulted in relatively large adherent liposomes, suggesting that PICK1 is prone to perform abscission-like scission of liposomes rather than tubulation as seen for other N-BAR proteins. This is in accordance with the punctate rather than tubular localization pattern of PICK1 in cells even upon overexpression.

Previous coarse grained MD simulations of the endophilin N-BAR domain on liposomes evolving to tubular networks (reticulation) showed early stages of deformation, termed budding, with endophilin lining up at the bottom of furrows to bulge out membrane buds an order of magnitude bigger than would fit the shape of the BAR domain (46). Notably, our recent Small Angle X-ray Scattering (SAXS) structure of PICK1 demonstrated elongated oligomers of PICK1 (35). Here we propose that this linear arrangement of PICK1 might evolve such initial buds into abscission events rather than tubular network.

On the other hand, TEM studies of liposome deformation by R197Q or R247H displayed tubular structures, while liposomes also incubated with R247H or R185Q formed small vesicular structures. Fission efficacy for N-BAR proteins was previously described as a balance between the fission promoting capacity of the amphipathic helices and fission restraining action of the oligomerized BAR domain, which stabilize tubular structures (17), although this view has been contested (34, 46). For PICK1, the oligomers support abscission-like fission, but the reduced hill coefficient for R185Q and R247H, suggest a compromised BAR-BAR oligomerization leading to more endophilin-like, tubular deformation and fission into small structures. Indeed, the coding variants (and in particular the two dominant negative mutations R158Q, and R247H) sit in a patch where they might alter the oligomerization interface according to our previously published SAXS structure (35). Unfortunately, we were unable to assess this directly by SAXS due to insufficient stability of the purified coding variants. The increased abscission efficacy provides nonetheless a credible mechanistic explanation for the dominant negative function of R158Q, and R247H.

A comparable set of disease-associated mutations in the BAR domain protein Bridging Integrator 1 (BIN1), D151N and R154Q, located slightly closer to the tip of the BAR domain, have previously been characterized in such a role (40). Similar to the PICK1 coding variants, the BIN1 mutations reduced membrane association without affecting curvature sensing. However, unlike our coding variants of PICK1, they compromise tubulation as a result of reduced oligomerization, highlighting that subtle perturbations to the oligomerization interface may dramatically alter the ability of N-BARs to shape lipid membranes. Ultimately, further insight into the membrane molding action of PICK1 and the changes imposed by the coding variants will rely on high-resolution structural data on lipid membranes.

The ability of PICK1 to deform lipid membranes in a cellular context was originally associated with budding of SGs from the TGN in growth hormone- and insulin-secreting cells (20, 21). Surprisingly, we did not see changes in proinsulin levels upon PICK1 KD as reported previously (20, 22) or after re-expression of the coding variants, suggesting that the function of PICK1 in INS-1E cells is likely more prominent after budding from the TGN. The subsequent process of SGs maturation entails removal of excess material such as syntaxin6, vesicle associated membrane protein 4 (VAMP4), furin, and both mannose 6-phosphate receptors (MPRs) from the SGs. This process relies on clathrin as the coat proteins as well as AP-1A and Golgi-localized, γ-ear-containing, ADP-ribosylation factor-binding proteins (GGAs) as adaptors recruited by ARF1 (12, 14, 47). Several lines of evidence presented here imply that the function of PICK1 extends to, not only the initial budding from TGN, but also the egress from the ISGs (see figure 6 and 7). Consistent with this idea, we recently demonstrated that molecular replacement of PICK1 with a fission incompetent PICK1 variant with mutations in the amphipathic helix in the N-BAR domain region (PICK1 V121E-L125E) resulted in larger insulin granules (24). Yet, the combined *in vitro* experiments and cellular imaging suggest that the functional effect of the coding variants identified in T2DM patients actually accelerated PICK1 egress from ISGs (see figure 8D). It is unclear how this may compromise insulin storage, but the tendency to produce more, small (≤70nm) PICK1 associated insulin granules suggest that insulin may to some extend follow egress if the process takes place prematurely e.g. prior to the pH and Ca^2+^ dependent condensation of insulin (see figure 8D).

In summary, four coding variants in the PICK1 BAR domain, which was discovered by WES in a group of Danish T2DM patients provided us with novel geno-phenotype relations, which are relevant for BAR domain proteins in general. Further, these mutations pointed us to a role of PICK1 in a novel trafficking pathway involved in maturation of insulin granules. We propose that this pathway might serve to regulate removal of insulin for e.g. lysosomal or autophagocytic degradation in response to changes in glucose concentration, but this requires further studies.

## Methods

### Cell cultures

COS7, a cell line derived from an African green monkey fibroblast, was cultured in Dulbecco’s Modified Eagle Medium (DMEM) 1885 containing 10% fetal bovine serum (FBS), 1% penicillin/streptomycin (P/S) and 1% L-glutamine, at 37°C in a humidified 10% CO_2_ atmosphere. INS-1E is an insulin-producing cell line derived from a rat β-cell. INS-1E cells were maintained in RPMI 1640 (with HEPES and 1% L-glutamine) or RPMI 1460 (with HEPES + Glutamax), both containing 10% FBS, 1% P/S, and 1.5% 100x RPMI supplement (1 mM Na-Pyruvate and 50 µM 2-mercaptoethanol) at 37°C in a humidified 5% CO_2_ atmosphere. The GRINCH cells are insulin-producing cells with stable expression of eGFP-proinsulin, a gift from Prof. Peter Arvan, University of Michigan Medical School, Michigan, USA (36). GRINCH cells were maintained in RPMI (with HEPES) including 10% heat-inactivated FBS, 1% P/S, 1mM Na-Pyruvate, 2 mM L-glutamine and 50 µM 2-mercaptoethanol, at 37°C in a humidified 5% CO_2_ atmosphere.

### *Drosophila* genetics

The fly lines carrying the UAS-driven *PICK1A*^*WT*^-*HA* and UAS-driven *GFP-Golgin245* transgenes (24) and the *PICK1*^*1*^ and *PICK1*^*2*^ deletion alleles (48) have been described previously. The *c929-Gal4* driver for expression in large high-capacity peptidergic neurons (49) was a gift from Dr. Paul Taghert, Washington University, Missouri, USA. The human missense mutation R247H corresponds to K249H in *Drosophila*, and The HA-tagged *PICK1A*^*K249H*^ transgene was generated in a similar way as UAS-PICK1A^WT^-HA and inserted into the same attP acceptor site on the 3^rd^ chromosome (M{3xP3-RFP.attP]ZH}-86Fb attP) using Phi31C recombination. In brief, the preexisting pUASTattB-PICK1A-HA construct was mutagenized by megaprimer mutagenesis to introduce mutations into the *PICK1A* coding region corresponding to the K249H alteration of the protein sequence. Embryo injections and transformant selection was performed by BestGene Inc. (CA, USA). Flies were reared on standard cornmeal food at 25°C. Genotypes used for immunostaining: WT PICK1A-HA rescue (*w*^*1118*^; *w+ c929-Gal4 PICK1*^*1*^ */PICK1*^*2*^; *w+ M*{*3xP3-RFP*.*attP*}*ZH-86Fb UAS-PICK1A-HA/w+ UAS-GFP-Golgin245*), PICK1A^K249H^-HA rescue (*w*^*1118*^; *w+ c929-Gal4 PICK1*^*1*^ */PICK1*^*2*^; *w+ M*{*3xP3-RFP*.*attP*}*ZH-86Fb UAS-PICK1A*^*K249H*^ *-HA/w+ UAS-GFP-Golgin245*).

### Molecular biology -DNA constructs

The pET41 vector encoding GST-PICK1 for bacterial expression (50)., and the peYFP C1 vector encoding YFP-PICK1 (28) have been described previously. Both vectors were edited using the QuickChange® kit (Stratagene) to generate constructs encoding PICK1 coding variants: R158Q, R185Q, R197Q, and R247H.

The FUGW vectors encoding shRNA targeting rPICK1 with expression of eGFP (KD) and eGFP-rPICK1 (KD + WT) were gifts from Dr. Robert Malenka, Stanford University, California, USA (30). The shRNA targeting rPICK1 was deleted to generate the ctrl vector as described (24). The PICK1 missense mutations were introduced into the KD + WT FUGW vector using the QuickChange® kit (Stratagene). The pHSynXW vector encoding eGFP-rPICK1 was a gift from Prof. Richard Huganir, Johns Hopkins University, Baltimore, USA. We replaced eGFP with mCherry using the QuickChange® kit. The myc-BioID2-PICK1 construct was purchased from Thermo Fisher Scientific and mPICK1 was fused to myc-BioID2 (39) with a linker region (GGGS).

### Lentiviral production

Lentiviral production was carried out as described previously (24). In short, HEK293T cells were cotransfected with the appropriate FUGW vectors and two helper plasmids (pBR8.91 and pMDG (PlasmidFactory)) using lipofectamine (Invitrogen, life technologies™) and 10 mM Na-Butyrate. The supernatant was collected and ultra-centrifuged at 20,000 rpm for 2 hours. The pellet was resuspended in MEM medium (gibco®, Thermo Fisher Scientific) and stored at −80°C.

### Protein expression and purification

*E. Coli* (BL21 DE3 pLysS) cultures were transformed with pET41 plasmids encoding either GST-PICK1 WT or the four PICK1 coding variants. Bacteria were pre-cultured in lysogeny (LB) medium (supplemented with kanamycin and chloramphenicol) and incubated on rotating shakers at 37° C overnight. Bacteria pre-cultures were transferred to 900 mL LB medium and grown at 37° C to ∼OD 0.6. Protein expression was induced with 1 mM Isopropyl β-D-1-thiogalactopyranoside and bacteria were incubated overnight at 20° C. Bacteria were harvested by centrifugation and resuspended in lysis buffer (50 mM Tris, 125 mM NaCl, 2 mM dithiothreitol (DTT) (Sigma-Aldrich), 1% Triton X-100 (Sigma-Aldrich), 20 µg/ml DNAse 1 and one tablet of complete protease inhibitor cocktail (Roche) per 100mL buffer). The resuspended pellet was frozen to −80° C. The bacterial suspension was thawed at 4° C and cleared by centrifugation (36.000 x g for 30 min at 4° C). The supernatant was collected and incubated with glutathione-sepharose 4B beads (GE-Healthcare) for 1.5 hours at 4° C on a rotator. The beads were pelleted at 4.000 x g for 5 min, the supernatant was discarded, and the beads were resuspended in wash buffer (50 mM Tris, 125 mM NaCl, 2 mM DTT and 0.01% Triton X-100). The washing step was repeated three times. The washed beads were transferred to Bio-Spin® chromatography columns (Bio-Rad Laboratories, Inc. cat. #7326008) and washed in 1mL wash buffer. Bead solution on the column was incubated with 3 µL 50U Thrombin (EMD Millipore, Novagen®) in 250 µL wash buffer overnight at 4° C on a rotator. PICK1 was eluded on ice and absorption at 280 nm was measured on a TECAN plate reader or a Thermo Scientific™ NanoDrop 2000c.

The protein concentration was calculated using lambert beers law (εA280PICK1=32320(cm*mol/L)-1). Alexa Fluor 488 C_5_ maleimide (Invitrogen™) labelling of GST-PICK1 WT and the coding variants was prepared as above, but without addition of DTT. Alexa Fluor 488 C_5_ maleimide was added to the Bio-Spin® chromatography columns prior to Thrombin restriction. The columns were sealed and incubated on a rotator for 4-16 hours at 4 ° C. The bead solution was washed 5x with wash buffer removing unconjugated dye. Thrombin was added and protein was eluded as described.

### Fluorescence polarization assay

Fluorescence polarization measurements were carried out in competition as previously described (50, 51), with a fixed concentration of PICK1-WT, PICK1 coding variants (non-saturating) and a fluorescent tracer (Oregon-Green DATC11, 20nM). The 15-residue peptide from the C-terminal region of the human dopamine transporter (RGEVRQFT-LRHWLKV) was purchased from Schafer-N A/S with >95% purity. The peptide was dissolved in wash buffer at a concentration of 2 mM, and a final assay concentration of maximally 1 mM was used. Following 15 mins of pre-incubation, increasing concentrations of unlabeled DATC15 peptide were added and incubated for an additional 20 mins on ice in a Corning® 96-well half area black flat bottom polystyrene nonbinding surface microplate. Fluorescence polarization was measured using an Omega POLARstar plate reader at 488 nm and 535 nm.

### Liposome preparation

We prepared unilamellar liposomes from bovine brain extract (Type I, Folch fraction I, Sigma-Aldrich) by following a standard hydration/extrusion procedure (18). The lipids were dissolved in 1 mL chloroform (0.5 mg or 1 mg for the liposome deformation assay or TEM imaging, respectively) before addition of 2% DiD (w/w), only for the liposome deformation assay. Lipids were dried while rotating using N_2_-gas, creating a thin lipid film, followed by vacuum dehydration for at least 10 hours to remove residual chloroform. Lipids were rehydrated in a sterile 200 nM D-Sorbitol solution (pH 7.4) to a final concentration of 0.5 or 1 mg/mL for the liposome deformation assay or TEM imaging, respectively. The rehydrated lipids were put through 8 “freeze-thaw” cycles; frozen using liquid nitrogen and thawed in a 40-50°C water bath. The liposomes were extruded through a 19 mm polycarbonate Whatman™ Nuclepore™ Track-Etched membrane with a pore size of 1 µm using a LiposoFast liposome extruder (Avestin). The liposomes were extruded 15 times before being diluted 1:200 in sterile phosphate buffered saline (PBS) with a total salt concentration of 100mM (94mM NaCl, 3.1mM Na2HPO_4_ and 0.9mM NaH2PO_4_, pH 7.4).

### Flow cytometry

We modified the previously described SLIC assay (32) to a high-throughput analysis by application of flow cytometry. Data was aquired on a LSRII, BD Biosciences (New Jersey, USA) flow cytometer and analysed using FlowLogic software (Inivai). Increasing concentrations of purified PICK1 WT and PICK1 coding variants labelled with Alexa Fluor™ 488 (AF488) (average labelling degree 30%, concentrations from 3 to 3000nM) were incubated with a fixed concentration of liposomes (∼2.5 µg/mL, 2% DiD (w/w)) and left at room temperature (RT) for 1 h prior to acquisition. Fluorescence intensities for DiD and AF488 were detected by photomultiplier tubes (PMTs) after passing through 660/20 nm and 505LP, 530/30 nm light filters, respectively. DiD was excited by a 633 nm JDS Uniphase HeNe laser (17 mW) and AF488 by a Coherent Sapphire 488 nm, air cooled laser (20 mW). Events were triggered by fluorescence, using the 488 nm laser filtered through a 685 LP filter, and 695/40 nm bandpass filter. The voltage applied to the PMT detecting events was set to 500V with a threshold value of 200, defined as the highest gain allowing a maximum of 1 event/s when running a filtered phosphate buffered saline solution. The voltages applied to the detectors of AF488 and DiD fluorescence were kept constant throughout experiments at 335V and 465V, respectively. We recorded at least 100,000 events per condition for each experiment, with a constant number of events per condition within each experiment.

For fission analysis, we applied gates removing >99.9% of the DiD background noise, acquired when running samples containing AF488 labelled protein diluted in PBS. The resulting data was applied in a kernel density estimation with 512 equally sized bins on a log scale ranging from 2 to 5.5 (A.U.) and normalized to the population obtained from a sample only containing liposomes by subtraction.

For binding and MCS analysis, we applied gates removing >99.9% of both DiD and AF488 background noise from events in samples containing just AF488 labelled protein and liposomes, respectively. The overall binding ability was estimated as the percentage of events surviving the gate. MCS was assessed as the slope of a linear regression model in a log-log graph of relative protein density (AF488/DiD) vs relative liposome size (√DiD).

### TEM imaging

Purified PICK1 WT and PICK1 coding variants were preincuabted with liposomes for 1 h at room temperature (RT), with final concentrations of 0.3 µM protein and 0.001 µg/mL liposomes. Grids were prepared by glow discharging for 30 seconds. 5 uL of protein-liposome mixture was added to the grids and incubated for 1 minute before 5 µL 2% uranyl acetate was added for an additional minute of incubation. The grids were washed with sterile H_2_O and filter paper was used to remove excess liquid. The grids were examined with a Philips CM 100 TEM at a voltage of 80kV, and the images were acquired with an Olympus veleta CCD camera.

### Transient transfection and transduction

A day prior to transfection, cells were seeded out on polyornithine-coated coverslips in 6-well tissue culture plates. Transfection was performed using Lipofectamine (Invitrogen, Thermo Fisher Scientific) and optiMEM (gibco®, Thermo Fisher Scientific). The protocol provided by the manufacturer was followed with a ratio of lipofectamine to DNA of 3:1. 0.5 and 1 µg DNA/mL was used for COS7 and INS-1E cells, respectively. Transfection-time was set to 5 hours or overnight where after optiMEM was replaced by culture medium. Experiments were carried out 48 hours post transfection. For live-cell microscopy cells were seeded on polyornithine-coated Lab-TEK II eight-well chambers at a density of 20,000 cells/well or on poly-L-lysine-coated MatTek microwell dishes (MatTeK Corporation) at a density of 60,000 cells/dish and grown for three days before experiments.

To optimize transduction efficiency, transductions were performed by spinoculation. 20 µL of the lentiviral suspension was added to 3×10^6^ INS-1E cells in 3 mL reheated medium, and centrifuged at 800 g, 32°C for 2 hours. The supernatant was aspirated and INS-1E cells were resuspended in preheated culture medium and transferred to 75T-tissue culture flasks (TPP®, Sigma-Aldrich). INS-1E cells were incubated for a minimum of 4 days to recover and initiate expression.

### Immunocytochemistry

#### Antibodies

Primary antibodies for immunostaining: mouse anti-clathrin (BD Biosciences, 610499) (1:500), chicken anti-GFP (Abcam, ab13970) (1:2000), rat anti-HA (Roche, clone 3F10) (200 pg/ml), chicken anti-PICK1 (Novus Biologicals, NBP1-42829) (1:500), mouse anti-PICK1 (custom-made, 2G10 clone) (1:500) (described in (48)), mouse-anti proinsulin (Abcam, ab82698) (1:1000), rabbit anti-insulin (Cell signaling, C27C9, #3014) (1:1000), guinea pig-anti insulin (Abcam, ab7842) (1:500), rabbit-anti TGN38 (Sigma-Aldrich, T9826) (1:2000), rabbit anti-syntaxin 6 (Synaptic Systems, 110-062) (1:2000). Secondary antibodies: Alexa Fluor® 488 conjugated goat anti-chicken (Abcam) (1:500), Alexa Fluor® 568 conjugated goat anti-mouse/rabbit (Thermo Fisher Scientific) (1:500), Alexa Fluor® 647 conjugated goat anti-mouse/rabbit (Thermo Fisher Scientific) (1:500) and Alexa Fluor® 647-conjugated goat anti-rat (Invitrogen, A21247) (1:500). For STED imaging following secondary antibodies were used: Abberior® STAR far red 640 goat anti-rabbit (Abberior GmbH) (1:500), Abberior® STAR orange 561 goat anti-mouse (Abberior GmbH) (1:500) and Alexa Fluor® 488 conjugated goat anti-guinea pig (Abcam, ab150185). The Abberior ® secondary antibodies were kindly provided by Abberior instruments.

Antibodies for dSTORM immunostaining: rabbit anti-insulin (Cell signaling, C27C9, #3014) (1:500), CF® 568 (Biotium) conjugated donkey anti-rabbit (Jackson ImmunoResearch Labs, 711-005-152) and Alexa Fluor® 647 (Invitrogen) primary conjugated chicken anti-GFP (Abcam, ab13970) or GFP nanobody (Chromotek, gt-250). Antibodies for immunoblotting: mouse anti-PICK1 (NeuroMab, Q9NRD5) (1:500), chicken anti-GFP (Abcam, ab13970) (1:1000), rabbit anti-chicken-HPR (Invitrogen, 31401) (1:10,000), goat anti-mouse-HRP (Invitrogen, 31430) (1:10,000). Loading control: HRP-conjugated anti-β-actin (Sigma-Aldrich, A3854) (1:20,000).

### Antibody conjugation

Secondary antibodies used for dSTORM were fluorescently labelled by NHS-ester fluorophore conjugation. 50µL antibody solution (donkey anti-rabbit, chicken anti-GFP or GFP nanobody) (1.2mg/mL) was added to 6 µL NaHCO_3_ (1M) and 1.25 µL NHS-ester fluorophore (CF®568 or Alexa Fluor® 647) (1mg/mL). The mixture was incubated for 2 hours at RT and antibodies were isolated using illustra™ NAP-5 columns (GE Healthcare Life Sciences™), resulting in an average labeling rate between 0.8 and 1.

### Immunostaining

Cells were seeded on polyornithine-coated coverslips in 6-well tissue culture plates with a density of 250,000 cells/well (COS7 cells) or 500,000 cells/well (INS-1E cells) four days prior to fixation. Staining of INS-1E cells for dSTORM imaging was prepared as described previously (24). For confocal and SIM imaging cells were washed in ice-cold PBS and fixed with 4% paraformaldehyde, 10 min on ice and 10 min at RT. COS7 cells were washed with PBS and milliQ, and mounted on a glass slides with Prolong® Gold antifade mounting reagent. INS-1E cells were washed in PBS and permeabilized for 30 min in PBS containing 0.2% saponin and 5% goat serum (GS). Subsequently, cells were incubated with primary antibodies (diluted in PBS with 5% GS) for 1 h at RT or overnight at 4°C and washed in PBS prior to incubation with secondary antibodies (diluted in PBS with 5% GS) for 30 min. Cells were washed in PBS and milliQ, and mounted on a glass slides using Prolong® Gold antifade mounting reagent. Glass slides were stored in the dark at 4°C. To visualize PICK1-HA and GFP-Golgin245 localization in *Drosophila* peptidergic somata, pharate adult stage pupae were dissected on Sylgard slabs in PBS and fixed in 4% formaldehyde for 30 min on ice. Washed 6 x 10 min in PBX (PBS + 0.3% Triton X-100), blocked in 5% GS in PBX for 30 min at RT, and incubated with primary antibody in blocking buffer over night at 4°C. This was followed by 6 x 10 min washes in PBX at RT and incubation with secondary antibody in PBX with 5% GS for 2 hours at RT. Finally, the samples were washed in 6 x 10 min in PBX, 2 x 5 min in PBS and mounted on glass slides using Prolong® Gold antifade mounting reagent. The intrinsic GFP fluorescence was visualized directly without immunolabelling.

### Cell lysates

For immunoblotting and ELISA, cells were washed in ice-cold PBS prior to lysis in 1xRIPA buffer (10 mM Tris-HCl, 150 mM NaCl, 1% Triton X-100, 1 mM EDTA, 1mM EGTA, 0.5% NP-40, pH 7.4) containing 1 mM PMSF and phosphatase inhibitor cocktail 3 (Sigma-Aldrich). Cells were scraped off, centrifuged at 11,000 g for 40 min at 4°C and the supernatant was stored at −20°C. The protein content was determined using the bicinchoninic acid assay kit (Pierce™, Thermo Fisher Scientific) to adjust the protein concentration for immunoblotting and to calculate relative proinsulin and insulin content.

### Bio-ID

6×10^6^ INS-1E cells were seeded out in tissue culture flasks (TPP®, Sigma-Aldrich) a day prior to transient transfection. Transfection was performed as mentioned above, 5 µg DNA was used per flask (in optiMEM) and 5 hours after transfection replaced by F10-DMEM media (Thermo Fisher Scientific). 48 hours post-transfection cells were washed with PBS and scraped off. The cell suspension was centrifuged at 1000 g for 5 min at 4°C, supernatant was aspirated and the pellet was snap frozen using liquid nitrogen. The pellet was resuspended in lysis buffer (50 mM Tris-HCl, 150 mM NaCl, 1% Triton X-100, 0.1% SDS, 0.5% sodium deoxycholate, pH 7.4), rotated for 1 h at 4°C and centrifuged at 13,000 g for 15 min at 4°C. Subsequently the supernatant was transferred to prewashed (in lysis buffer) sepharose 4B beads (sepharose 4B beads (Sigma-Aldrich). The sepharose 4B bead suspension was incubated on a rotator for 1 h at 4°C and centrifuged at 1000 g for 5 min at 4°C. The supernatant was transferred to prewashed (in lysis buffer) magnetic streptavidin-coupled Dynabeads (Thermo Fisher Scientific) and incubated for 1 h at 4°C on a rotator. The supernatant was removed from the Dynabeads using a magnetic DynaMag rack (Invitrogen, Thermo Fisher Scientific). The Dynabeads were washed twice in low salt wash buffer (20mM Tris-HCl (pH 7.4), 100 mM KCl, 0.1% Triton X-100), twice in high salt wash buffer (20 mM Tris-HCl (pH 7.4), 500 mM KCl, 0.1% Triton X-100), twice in low salt wash buffer and once in normal wash buffer (20mM Tris-HCl (pH 7.4), 150 mM KCl, 0.1% Triton X-100), before resuspention in elution buffer (50mM Tris-HCl, 150 mM NaCl, 1% SDS, 1mM biotin)). The lysate was mixed with 5x Sodium Dodecyl Polyacrylamide (SDS-PAGE) loading buffer (0.75M Tris-HCl, 50% glycerol, 2.5% SDS, 10% β-mercaptoethanol, 0.2% bromophenol blue and 100 mM DTT) and subsequently analyzed by immunoblotting.

### Immunoblotting

Cells were seeded on polyornithine-coated coverslips at a density of 500,000 cells/well three days prior to lysis. 20 µg total protein lysate was mixed with 5xSDS-PAGE loading buffer, boiled at 90°C for 10 min loaded into an Any-kD precast gel (Mini-ProTEAN® TGX™) and run at 100V for 1 h. The size-separated proteins were transferred to polyvinylidene difluoride membranes (BIO-RAD) for 2 hours at 18 V and blocked for 1 h in 5% milk in wash buffer (PBS with 0.1% Tween-20). Membranes were incubated with primary antibodies for 1 h at RT or overnight at 4°C, washed and incubated with HRP-conjugated secondary antibodies for 30 min. Membranes were washed before developed using either the SuperSignal ELISA Femto Substrate (Thermo Fisher Scientific) or ELC Prime Western Blotting system (Sigma-Aldrich) and captured with a cooled CCD camera.

### Proinsulin and insulin ELISA

High range rat proinsulin and insulin ELISA kit immunoassays (ALPCO™) were used to measure the proinsulin and insulin content in INS-1E lysates. Cells were seeded at a density of 250,000 cells/well in twelve-well tissue culture plates four days prior to cell lysis. Protocol provided by the manufacturer was followed. Briefly, the kit consists of a 96-well microplate pre-coated with a monoclonal antibody for proinsulin or insulin. Lysates and HRP-conjugated secondary antibodies were added. Absorption was measured using an Omega POLARstar plate reader at 450nm and 590nm.

### Light microscopy

Confocal microscopy was performed on fixed coverslips using an inverted laser-scanning microscope (LSM) 510 or an upright LSM 710 (Carl Zeiss, Oberkochen, Germany) both with 63x/1.4 numerical aperture (NA) oil immersion apochromat objectives.

LSM 510 settings; Alexa Fluor® 488 and YFP fluorescent signals were detected using a 488 nm argon laser. Alexa Fluor® 568 and Alexa Fluor® 647 fluorescent signals were detected using a 543 nm helium-neon laser and a 633 nm helium-neon laser. LSM 710 settings; Alexa Fluor® 488, Alexa flour® 568 and Alexa Fluor® 647 fluorescent signals were detected using a 488 nm argon laser, a solid state 561 nm laser and a helium-neon 633nm laser, respectively. Channels were imaged separately. Confocal imaging of INS-1E cells was performed blinded.

Live-cell images of GRINCH cells were acquired on an inverted Nikon A1+ point scanning confocal microscope (Nikon, Japan) with 405, 488, 561 and 647 nm lasers and a 60x/1.4 NA oil immersion Apochromat objective. Cells were kept in a Krebs Ringer solution (20 mM HEPES (pH 7.4), 119 mM NaCl, 4.75 mM KCl, 2.54 mM CaCl_2_, 1.2 mM MgSO_4_, 1.18 mM KH_2_PO_4_, 5 mM NaHCO_3_) supplemented with 0.2% BSA and 11 mM glucose, in a heated and humidified chamber. Z-stack live imaging was performed with a total coverage of 0.500 µm in the z-plane. Recorded for 2-4 min per session. The eGFP and mCherry fluorescent signals were detected using the 488 nm and 568 nm laser, respectively.

### Super-resolution microscopy

dSTORM images were acquired on an ECLIPSE Ti-E epifluorescence/TIRF microscope (Nikon, Japan) with 405, 488, 561 and 647nm lasers (Coherent, California, USA) and a 100x/1.49 NA oil immersion Apochromat objective. Settings and conditions for dSTORM imaging are the same as previously described (24). dSTORM imaging on fixed coverslips was performed in a buffer containing β-mercaptoehtanol and an enzymatic oxygen scavenger system (10% (w/v) glucose, 1% (v/v) β-mercaptoethanol, 50 mM Tris-HCl (pH 8), 10 mM NaCl, 34 µg ml^−1^ catalase, 28 µg ml^−1^ glucose oxidase). Images were acquired at 10,000-30.000 cycles of one frame of 561 nm laser activation followed by one frame of 647 nm laser activation at 16 Hz per cycle. Shutters regulated the light path in order to separate the light between the frames. The 561 nm and 647 nm lasers were held constant at 0.6 and 1.1 kW cm^−2^, respectively, while a 405 nm laser was gradually increased to <0.1 kW cm^-2^.

SIM was performed on RT fixed coverslips using an Elyra PS.1 microscope (Carl Zeiss, Oberkochen, Germany) with 488, 561, and 642 nm lasers (Coherent, California, USA) and a 63x/1.4 NA oil immersion Apochromat objective. Z-stack images were acquired from top to bottom of cells (0.110 µm per interval) and channels were imaged separately.

STED images were acquired on fixed coverslips using a STEDYCON with 488, 561 and 640 nm lasers together with a 775 nm STED laser (all pulsed) (Abberior Instruments GmbH, Göttingen, Germany). The STEDYCON was attached to a Zeiss Axio imager/observer xx microscope with a 100x/1.46 NA oil immersion Apochromat objective. The 488 laser was used to obtain confocal resolution in cells stained for guinea pig anti-insulin and subsequently with the secondary Alexa Fluor® 488 conjugated goat anti-guinea pig antibody.

### Confocal image analysis

Image processing of confocal images was primarily performed using the ImageJ software program (Rasband W. S., ImageJ, U.S. National Institutes of Health, Bethesda, MD, USA). Cells in confocal images were outlined by region of interest (ROIs) in the 488-channel (YFP/GFP), in combination with the PICK1-channel to identify transfected/transduced cells. Prior to quantification of the immunosignal, background noise was subtracted and a threshold was set for each channel. These settings were held constant throughout each individual experiment. The images were converted to binary images and multiplied into each ROI of the original image, resulting in a total intensity value. To analyze the clusters, an analyze particle filter (available in ImageJ) was added with a minimum of 5 pixels per cluster and with a circularity of 0.10-1.00 before multiplying the binary image with the original image. For total immunosignal analysis, the immunosignals from the transfected/transduced cells within each image were normalized to the untransfected/untransduced cells. Quantification of the PICK1 and insulin immunosignal was performed blinded. Plotting of the PICK1 and insulin immunosignal per cell, shown with linear regression, excluded the PICK1 and insulin immunosignals above the ctrl cells, from each individual experiment.

To examine colocalization between PICK1 and different cellular markers, Van Steensel’s cross-correlation function was used on the processed images with the JaCoP plugin in ImageJ (52). The Van Steensel’s cross-correlation calculates the Pearson cross-correlation as the signal from channel one (ex. blue PICK1) shift relative to the signal from channel two (ex. Magenta TGN38) in the x-direction pixel per pixel. The peak at Δx = 0 indicates a specific colocalization and a value of 1 indicates total overlap. Analysis of the colocalization was performed blinded.

Analysis of YFP-Δ1-101PICK1 clusters in COS7 was performed using the Igor Pro version 6.34A software (WaveMertrics, USA). Clusters were separated by ROIs and chosen on the premises of a minimum area of 5 pixels, maximum deviation of 75%, fluctuation factor 1 and minimum ellipticity of 0.5. Background threshold was determined for each image due to high variation in cluster intensity. For presentation, the images are shown with an inverted gray scale.

### dSTORM image analysis

Localizations were fitted to the dSTORM movies with the ThunderSTORM plugin for imageJ (53). Images were filtered with the Wavelet filter (B-Spline), and localizations detected with the local maximum detector with a threshold of 2.0*std (Wave.F1). Drift was corrected with cross correlation and localization with uncertainty greater than 20 nm was filtered out. For the colocalization analysis, we used coordinate based colocalization (CBC) (54) as previously described for PICK1 and insulin clusters (24). The CBC was completed in 20 steps of 5nm, and the clusters were identified with the DBSCAN algorithm through the sklearn library in python 3.6, where a localization was classified as clustered if it had 10 localizations or more within a 40nm radius. Each localization within a PICK1 cluster was assigned a CBC value ranging from 1 to –1 based on the distance to clusters of insulin and vice versa. Values of 1 defines a perfect overlap between a PICK1 and an insulin cluster, whereas a value of –1 indicates no overlap between the clusters. DBSCAN was performed in 3D. Diameter was determined from the 2D area of the convex hull of the cluster in the X-Y plane by fitting the area to a circle. The software programs ImageJ and MATLAB (MathWorkers, Natick, MA, USA) were used for image visualization.

### SIM image analysis

Images were reconstructed using a built-in algorithm in the Zen software (Carl Zeiss, Oberkochen, Germany). A noise filter was added and held constant for each individual experiment, although always set between −4.2 and −4.5. Drift was corrected for with a bead alignment matrix. For 3D visualization we used the Amira software 2019.1 (Thermo Fisher Scientific, Waltham, MA, USA).

## QUANTIFICATION AND STATISTICAL ANALYSIS

Data was transferred to the GraphPad Prism software (GraphPad Software, San Diego, CA, USA) for statistical analysis and presentation. Data were tested for normality using either the D’Agostino-Pearson normality test or the Shapiro-Wilk normality test. For data with normality we used one-way ANOVA followed by either Dunnett’s or Turkey’s multiple comparisons test for statistical comparison, while data that were not normally distributed was tested for statistical comparison using the non-parametric Kruskal-Wallis test followed by Dunn’s multiple comparisons test. Two-way ANOVA followed by Dunnett’s multiple comparisons test was used for analysis of fractional cluster size distribution. Statistical significance for cumulative distributions was compared using the Kolmogorov-Smirnov test in MATLAB. Data is presented as mean ± SEM if nothing else is stated. P values of <0.05 were considered significant. n represents the number of cells or individual experiments performed. The built-in GraphPad analysis “identify outliers” was used for the dataset in Figure 1.

## DATA AND CODE AVAILABILITY

The codes for dSTORM image analysis generated during this study are available at [name of repository] [accession code/web link].

## ACKNOWLEDGEMENTS

We thank Prof. Torben Hansen, University of Copenhagen, Copenhagen N, Denmark and the Lundbeck Foundation Centre for Applied Medical Genomics in Personalised Disease Prediction, Prevention, and Care (LuCamp) (Denmark) for access to the WES data. We thank Nabeela Khadim and Anders Bohl Pedersen for excellent technical assistance. We also thank Thomas Hartig Braunstein and Pablo Hernandez-Varas from the Core Facility for Integrated Microscopy, Department of Biomedical Sciences, University of Copenhagen for help with microscopy, and to Elle Kielar Grevstad from the Biochemistry Optical Core, University of Madison-Wisconsin for help with live-cell microscopy. We thank Dr. Robert C. Malenka, University of Stanford, California, USA for the shRNA constructs, Prof. Richard Huganir, Johns Hopkins University, Baltimore, USA for the the pHsSynXW vector and Prof. Peter Arvan, University of Michigan Medical School, Michigan, USA for the GRINCH cells.

## AUTHOR CONTRIBUTIONS

*In vitro* work was conducted by R.C.A, N.R.C, J.H.S, M.S, J.R and G.N.H. Cellular experiments were performed by R.C.A with help from M.D.L, D.S, J.R and A.M.J. R.C.A and V.K.L performed the drosophila experiments. R.C.A, M.D.L, J.H.S, N.R.C, R.J, V.K.L, A.M.J, O.K, A.D.A, K.L.M and U.G designed the experiments. Data analysis was performed by R.C.A, M.D.L, J.H.S, N.R.C, V.K.L, J.R, M.P.K, R.H, O.K, A.D.A, B.H, K.L.M and U.G. The manuscript was written by R.C.A, K.L.M and U.G.

---

**Supplemental Figure 1 (Related to Figure 2).**
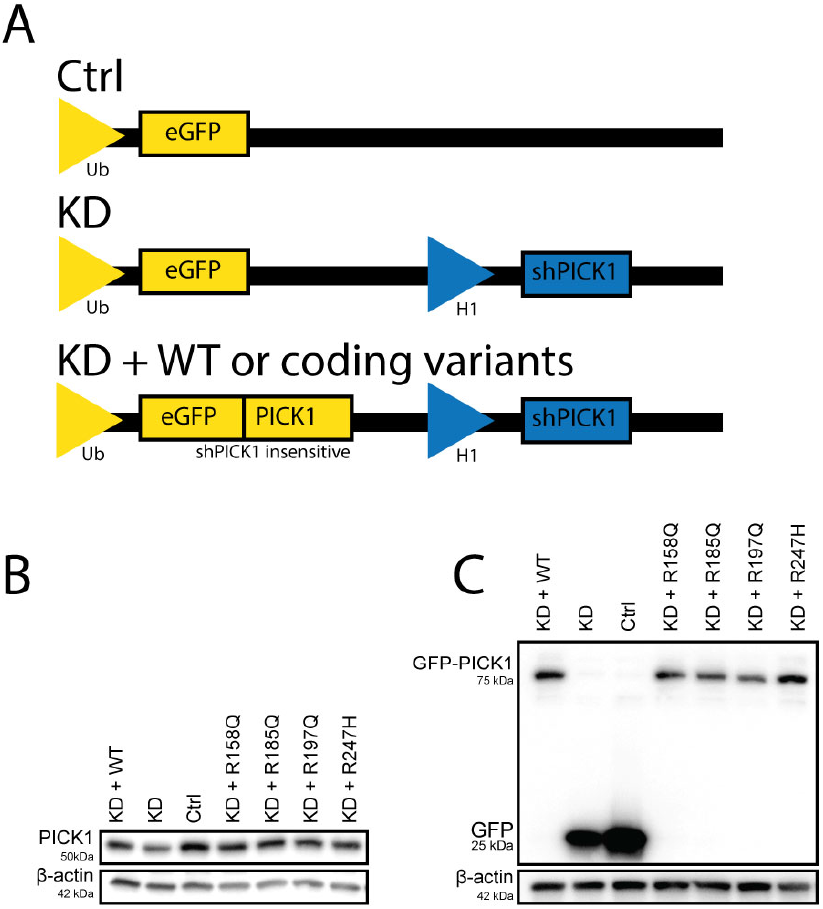
Immunoblotting show a reduced level of endogenous PICK1 with an expression of GFP or GFP-PICK1 in INS-1E cells transduced with the lentiviral constructs. (A) Schematic diagram of the different FUGW constructs; ctrl, KD, and KD + WT or the human mutations. (B) Immunoblotting of INS-1E lysates shows expression of endogenous PICK1, and (C) GFP(-PICK1). Note that low transduction efficiency result in only weak knock down as assessed in bulk by western blotting. Knock down in individual transduced cells as assessed by immunocytochemistry was much more robust as seen in figure 2, A-B.

**Supplemental Figure 2 (Related to Figure 2).**
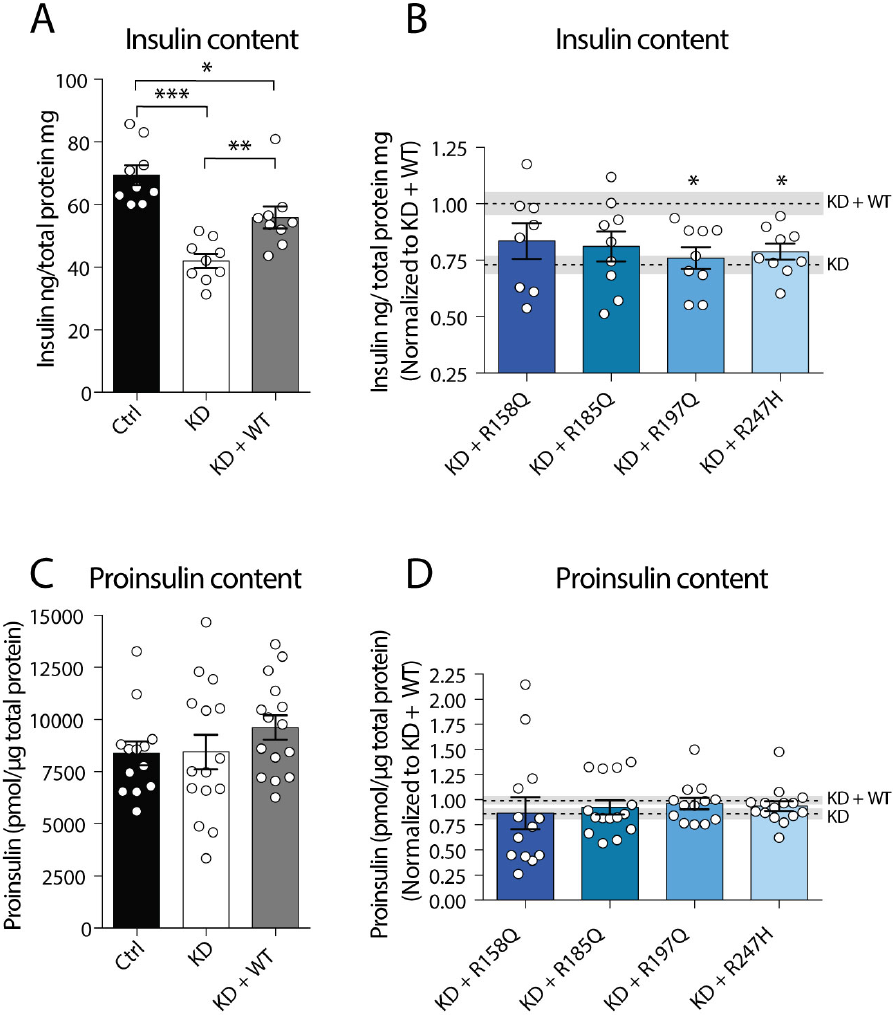
R197Q and R247H compromise PICK1 function in insulin storage but not proinsulin content in INS-1E cells as assessed in bulk by ELISA assay. Sandwich ELISA was performed on lysates from lentivirally transduced INS-1E cells. Bars are mean ± SEM with one-way ANOVA followed by Tukey’s or Dunnett’s multiple comparisons test if nothing else is stated. (A) Quantification of the insulin content in INS-1E cells expressing ctrl, KD and KD + WT, n = 8-9 samples pr. genotype. (B) Quantification of the insulin content from INS-1E cells expressing the four PICK1 coding variants, normalized to the mean insulin content from KD + WT from (A). The two dotted lines represent the mean insulin content ± SEM from KD and KD + WT from (A). n = 8-9 samples pr. genotype. (C) Quantification of the proinsulin content from INS-1E cells expressing ctrl, KD or KD + WT, n = 13-15 samples pr. genotype. (D) Quantification of the proinsulin content from INS-1E cells expressing the four PICK1 coding variants, normalized to the mean proinsulin content from KD + WT from (C). Data are shown as mean ± SEM with Kruskal Wallis test. The two dotted lines represent the mean proinsulin content ± SEM from KD and KD + WT from (C), n = 13-15 samples pr. genotype. (\**p*<0.05, \*\**p*<0.01, \*\*\**p*<0.001).

**Supplemental Figure 3 (Related to Figure 2).**
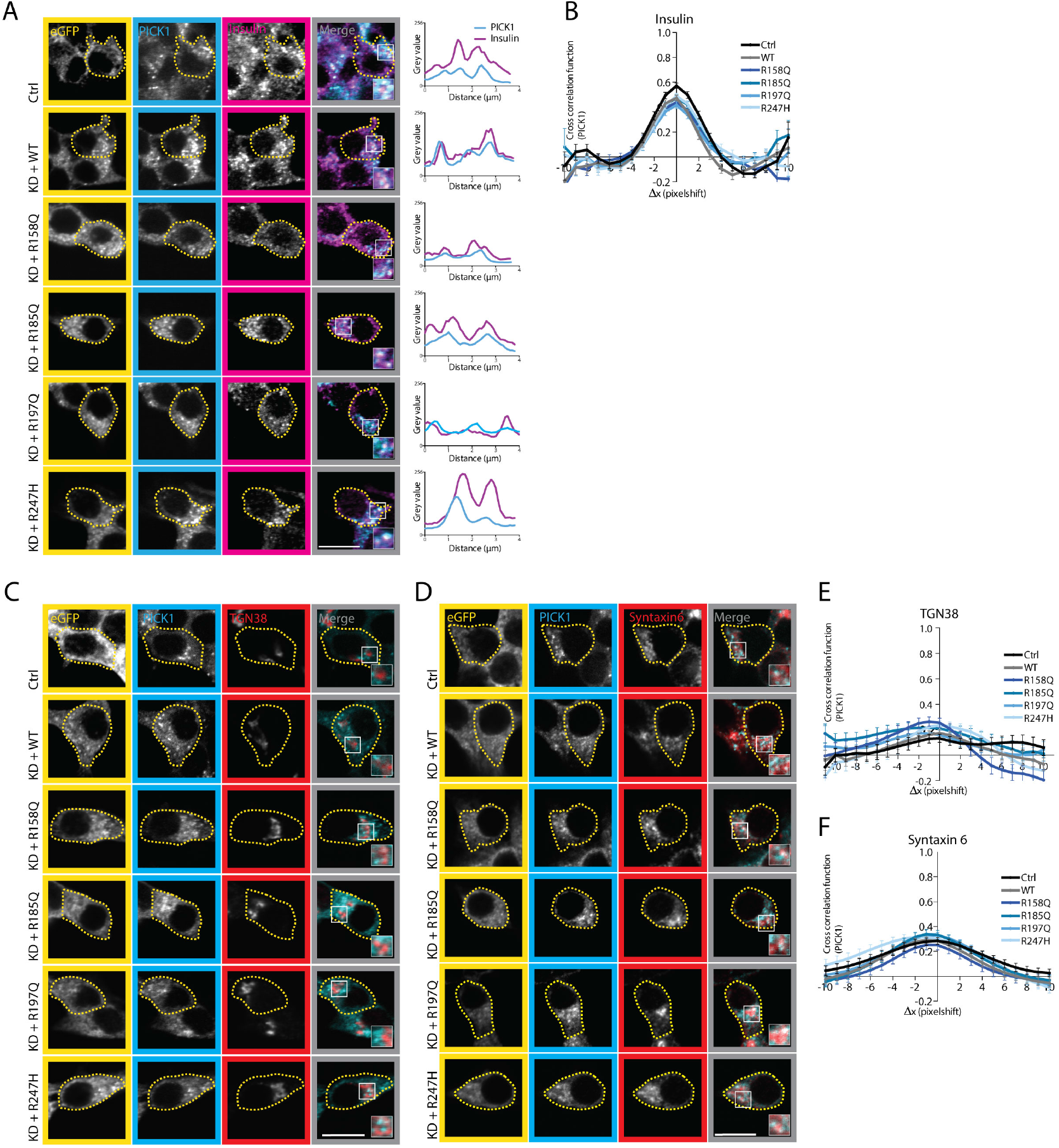
No change in the localization of PICK1 in the RSP with the four PICK1 coding variants. Representative confocal images of INS-1E cells transduced with the lentiviral constructs. Transduced cells (GFP-Positive) are outlined with a yellow dotted line. Merged images are with insets highlighting areas of overlapping localization between PICK1 and RSP markers. Zooms are gain adjusted for visualization. Scale bar = 10 µm. The colocalization was quantified by the Van Steensel’s cross-correlation function. (A) Representative confocal images of INS-1E cells immunostained for GFP (yellow), PICK1 (cyan) and insulin (magenta). *Right* profile plots from the insets. (B) Quantification of colocalization between PICK1 and insulin, ctrl (endogenous PICK1) (n =69), KD + WT (n = 64), KD + R158Q (n = 54), KD + R185Q (n = 64), KD + R197Q (n = 57), and KD + R247H (n = 58). (C) Representative confocal images of INS-1E cells immunostained for GFP (yellow), PICK1 (cyan) and TGN38 (a marker for the TGN) (red). (D) Representative confocal images of INS-1E cells immunostained for GFP (yellow), PICK1 (cyan) and syntaxin 6 (a marker for the TGN and early ISGs) (red). (E) Quantification of colocalization between PICK1 and TGN38, KD+ R158Q (n = 31), KD + R185Q (n = 31), KD + R197Q (n = 27), KD + R247H (n = 20), ctrl (n = 27) and KD + WT (n = 39). (F) Quantification of colocalization between PICK1 and syntaxin 6, KD+ R158Q (n = 26), KD + R185Q (n = 34), KD + R197Q (n = 28), KD + R247H (n = 27), ctrl (n = 37) and KD + WT (n = 34). Data are shown as mean ± SEM.

**Supplemental Figure 4 (Related to Figure 4).**
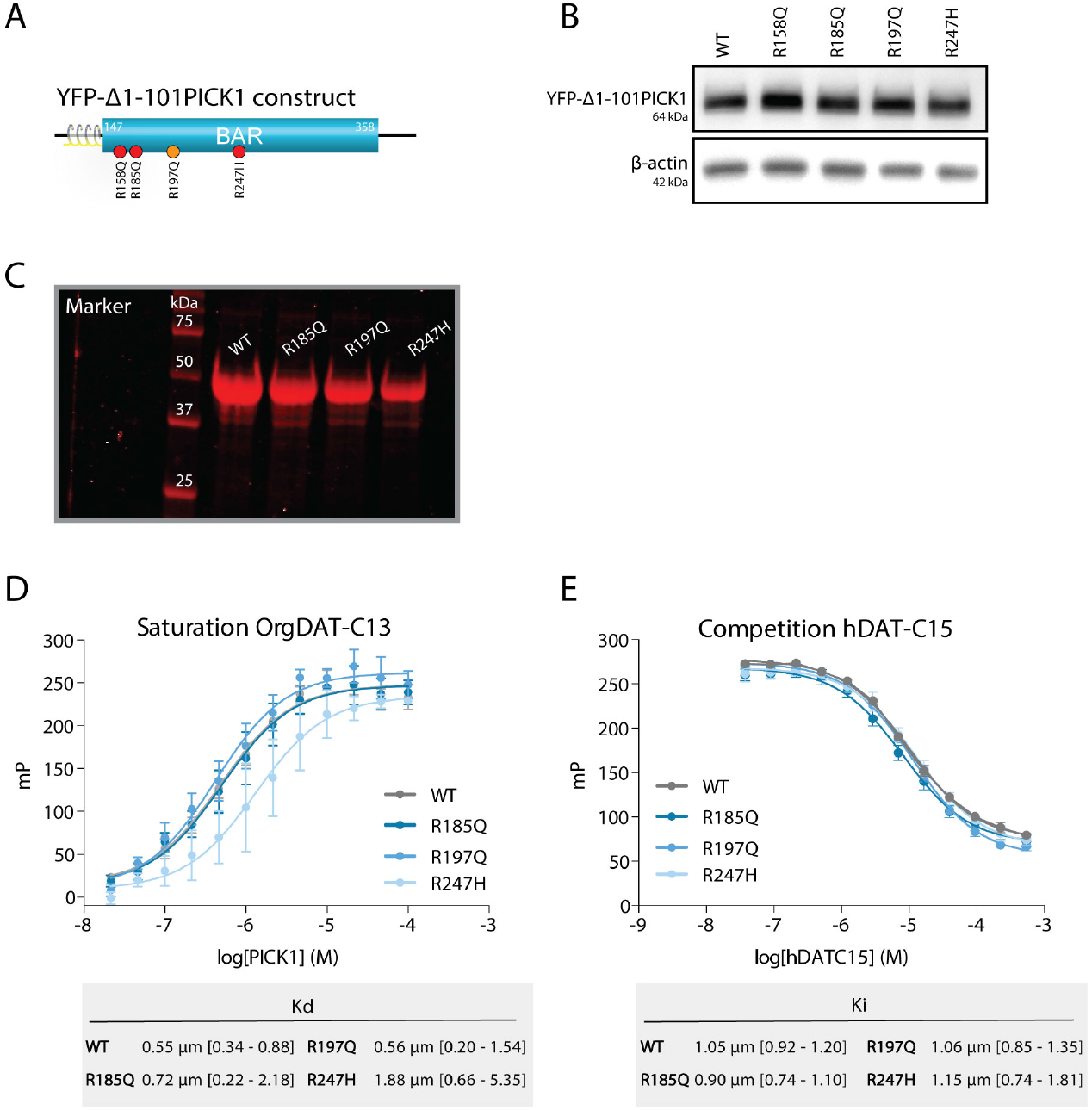
Expression and integrity of constructs used for Figure 4 and 5. (A) Schematic representation of the YFP-Δ1-101PICK1 construct comprising the amphipathic helix (H0) and BAR domain. (B) Immunoblotting shows the expression level of YFP-Δ1-101PICK1 WT, R158Q, R185Q, 197Q and Q247H in transiently transfected COS7 cells. (C) Representative SDS-page shows the expression level of purified full-length PICK1 WT, R185Q, R197Q and R247H, respectively. R158Q could not be purified. (D) Binding of the OregonGreen-DATC11 via the PICK1 PDZ domain to of purified PICK1 WT, R185Q, R197Q and R247H determined by fluorescent polarization (FP) saturation binding with affinities (K_D_) listed below. Data represented for n = 3 ± SEM. (E) FP competition binding of PICK1 WT, R185Q, R197Q and R247H competing for binding to DATC15 against OrG-DATC11, with affinity (K_i_) listed below. n = 6, data shown as mean ± SEM. The preserved PDZ domain binding test to the overall integrity of the purified proteins.

**Supplemental Figure 5 (Related to Figure 4 and 5).**
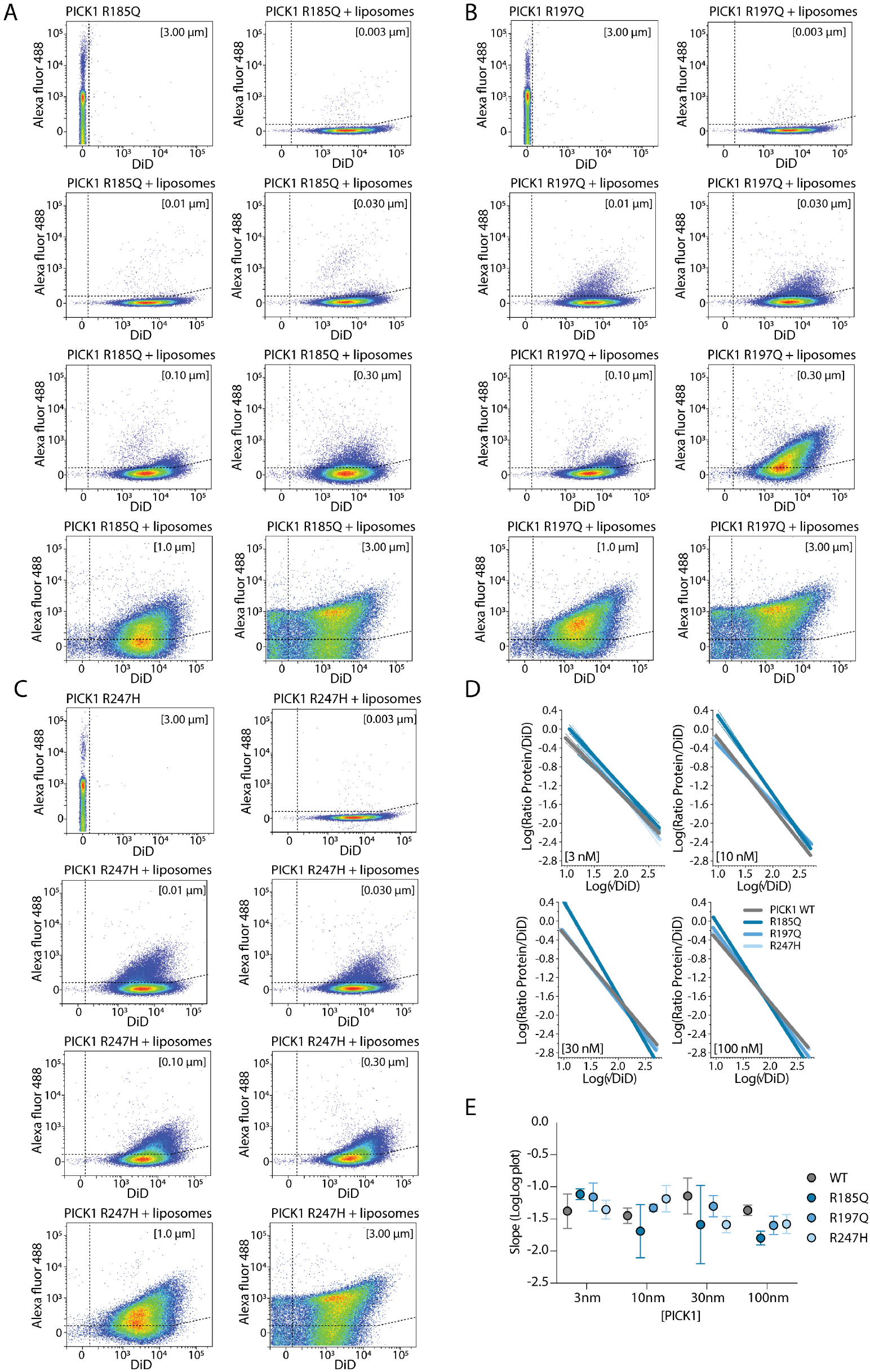
Flow cytometry plot of the purified AF488 labelled PICK1 coding variants binding to DiD labelled liposomes. (A-C) Representative two-parameter density plot of primary data output from flow cytometry showing fluorescent intensities (A.U.) of AF488 (PICK1) vs. DiD (liposomes) for samples containing liposomes or PICK1 alone as well as liposomes with increasing concentration of PICK1 WT (3 to 3000nM). Densities of observations indicated from low (blue) to high (red). (D) Log-log graph of protein densities of liposomes ((AF488/DiD)/ÖDiD) for a concentration range (3nM to 100nM – the concentration for which we observed no significant deformation) yielding MCS of PICK1 WT and coding variants to liposomes after 1 h incubation. The MCS is the slope of a linear regression fit of the relation between protein density (Ratio Protein/DiD) and liposome size in arbitrary units (√DiD). Pooled data from three experiments (E) Quantification of mean MCS from (D) as a function of concentration. Each dot represents the MCS for a given concentration of PICK1 WT or coding variant. Error bars indicate 95% confidence intervals of linear regression fits. No statistically significant difference between concentration or genotype (two-way ANOVA).

**Supplemental Figure 6 (Related to Figure 5).**
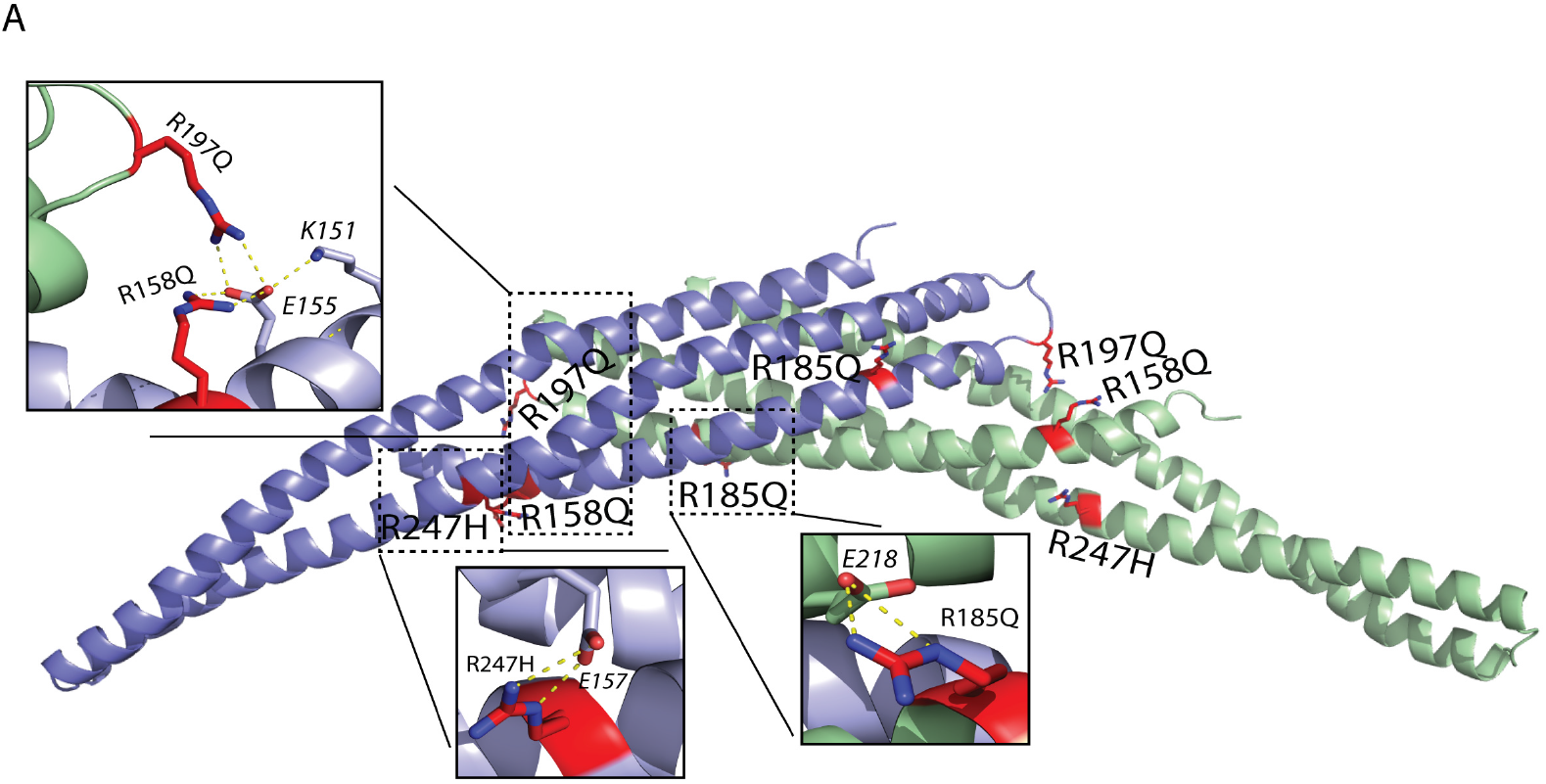
Structure of the dimeric PICK1 BAR domain with the position of four coding variants. (A) Ribbon diagram of the PICK1 BAR domain (threated based on its homology to arfaptin 2) shows the localization of the four identified coding variants in the homodimeric complex of PICK1, shown from the side (individual monomers represented as green and purple). *Bottom and top* show highlights of the PICK1 coding variants amino acid residues as sticks with nitrogen atoms (blue) and oxygen atoms (red). Putative charge-charge interactions with nearby negatively charged residues are shown with yellow dashed lines.

**Supplemental Figure 7 (Related to Figure 6).**
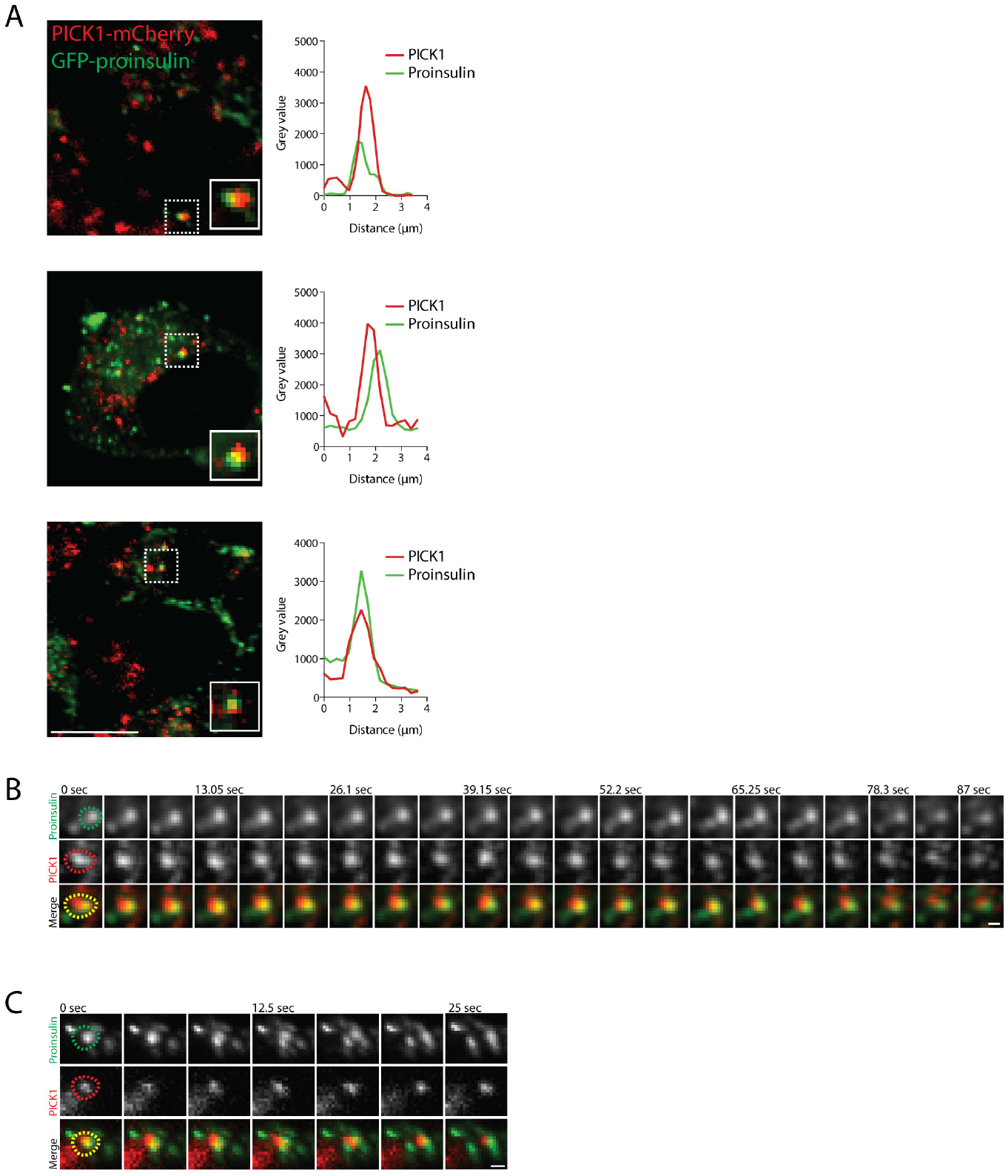
PICK1-positive clusters interact with proinsulin clusters. GRINCH cells were transiently transfected with PICK1-mCherry. (A) Merged images of PICK1-mCherry (red) and GFP-proinsulin (green) with insets highlighting areas of overlapping localization, scale bar = 10 µm. *Right* shows profile plots from the insets. (B and C) Rep-resentative merged z-stack (500nm confocal slice) images of PICK1 clusters (red) and proinsulin clusters (green) overlapping during live microscopy under steady-state conditions (11mM glucose). Time-lapse imaging was of two-color z-stacks, acquired every 4.34 seconds. Upper two rows represent a PICK1-mCherry positive cluster and a GFP-proinsulin cluster, respectively, both shown in greyscale. The third row shows merged images. Time is in seconds and scale bar = 1 µm.

**Supplemental Figure 8 (Related to Figure 6).**
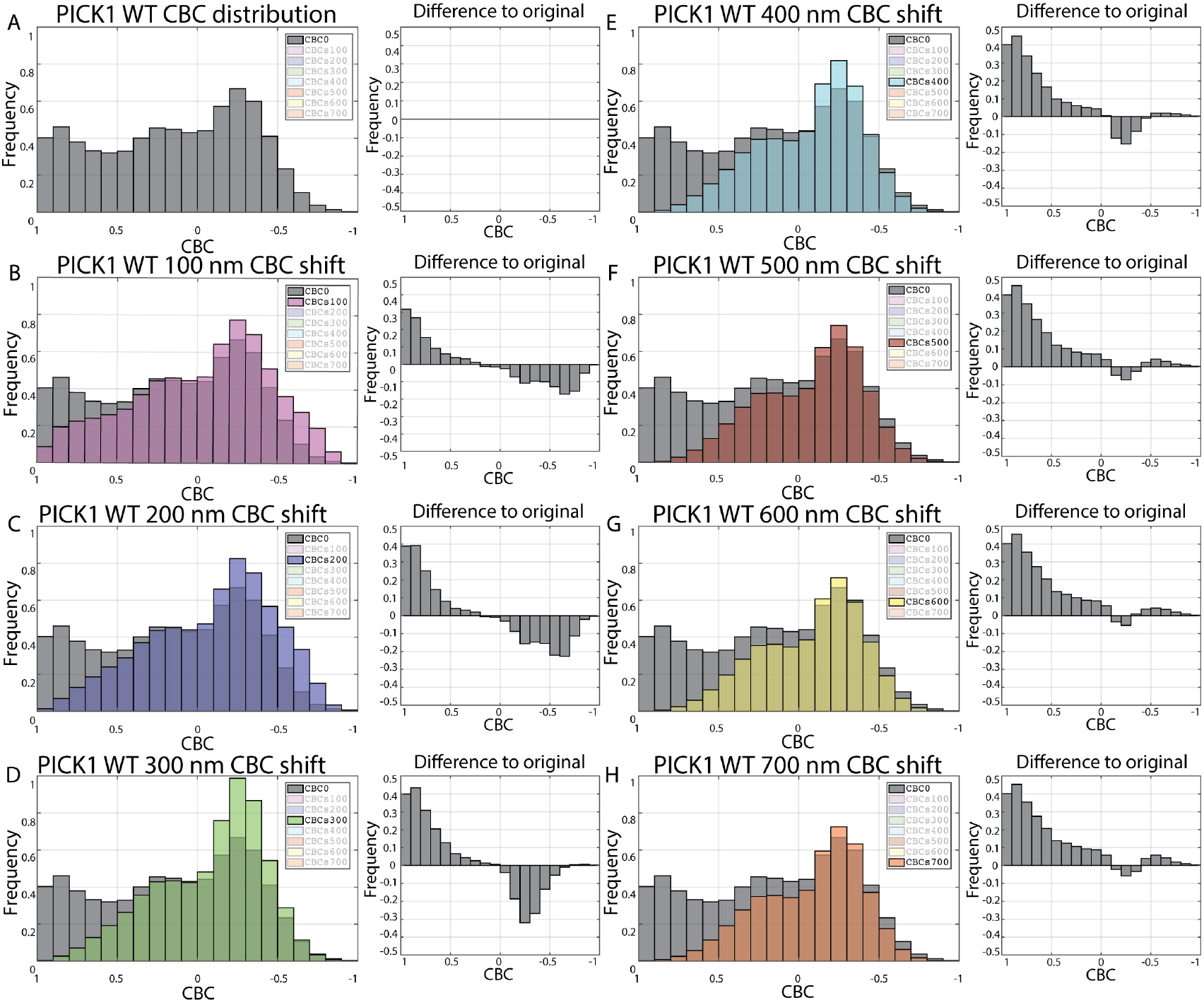
Insulin CBC shift analysis to eliminate random colocalization. (A) The CBC distribution of PICK1 localization to insulin granules in INS-1E cells transduced with KD + WT. (B-H) The PICK1 clusters were shifted +100 nm per interval in the x direction, and the CBC distribution of of PICK1 localization to the insulin granules are recalculated and compared with the CBC distribution from (A) (grey). *Right* panels show the difference in CBC between (A) and (B-H), which we refer to as random proximity subtracted CBC (rpsCBC) distribution. The analysis represents 3 analyzed images. Note that many points are not assigned a CBC values (NA) when shifted and this proportion increases with distance of the shift. The ‘difference to original’ plot show no further development beyond 500 nm.

**Supplemental Figure 9 (Related to Figure 7).**
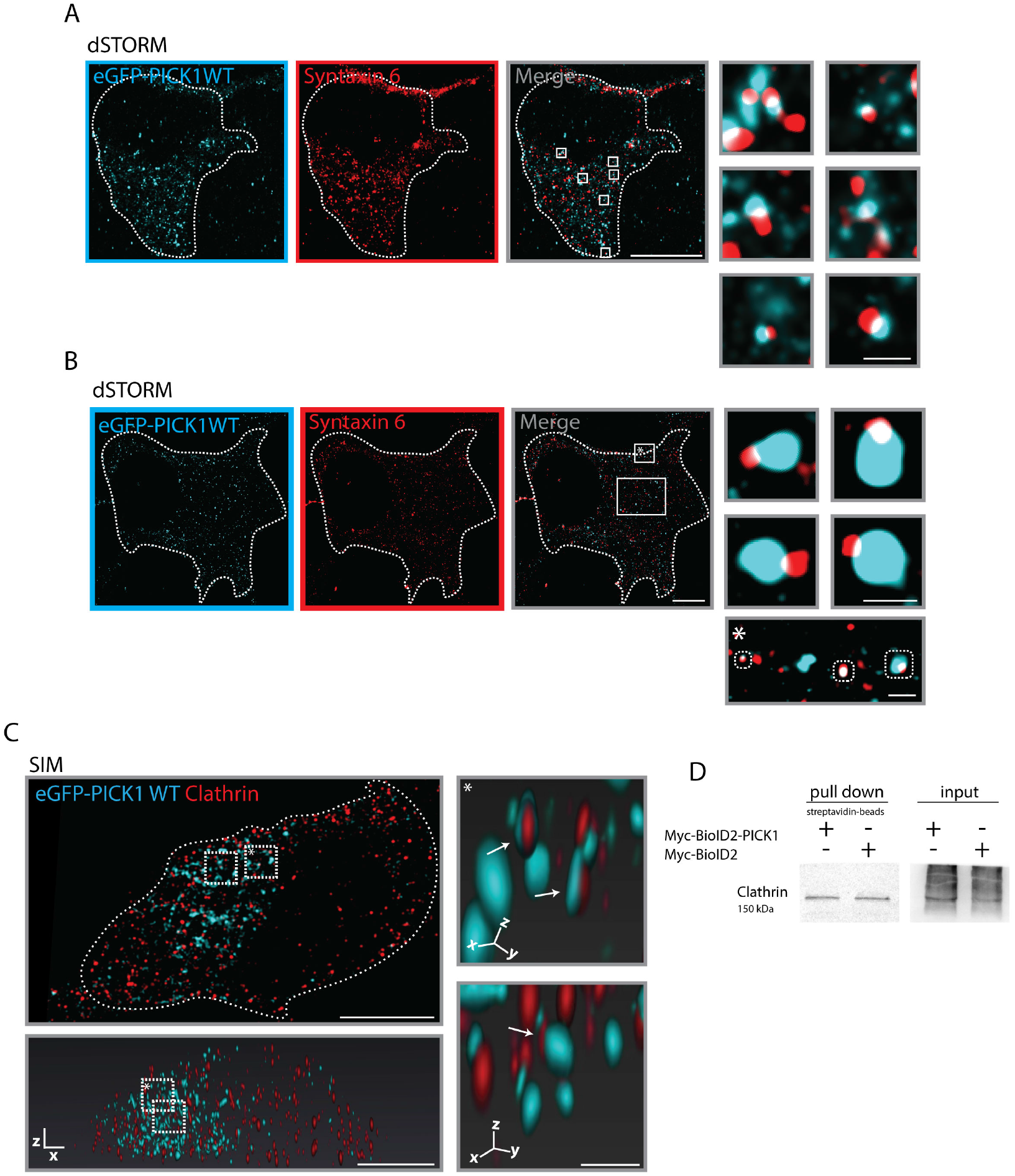
Syntaxin 6 and clathrin colocalization with PICK1. (A-B) Representative dSTORM images of INS-1E cells transduced with KD + WT, and immunostained for eGFP-PICK1 (cyan) and syntaxin 6 (red), scale bar = 5 µm. (A) Same INS-1E cell presented in figure 7D. *Right* shows insert with higher magnification of small overlapping PICK1 and syntaxin 6 clusters (<100 nm). Scale bar = 250 nm. (B) *Right* shows inserts with higher magnification of small syntaxin 6 vesicles overlapping with medium sized PICK1 clusters (∼150-200 nm). *Bottom* shows three different sizes of overlapping PICK1 and syntaxin 6 vesicles. Scale bar = 250 nm. (C) Representative SIM image of INS-1E cells transduced with KD + WT, and immunostained for GFP-PICK1 (cyan) and clathrin (red). Scale bar = 5 µm. *Bottom* shows same INS-1E cell in 3D reconstruction, and *right* shows insert with higher magnification of overlapping PICK1 and clathrin clusters (arrow). Scale bar = 500 nm. (D) INS-1E cells were transiently transfected with a Myc-BioID2-PICK1 construct. For control, INS-1E cells were transiently transfected with a Myc-BioID2 construct. Biotinylated proteins were pulled down from cell lysates with streptavidin-beads for immunoblotting against clathrin. n = 4 individual experiments.

**Supplemental Figure 10 (Related to Figure 8).**
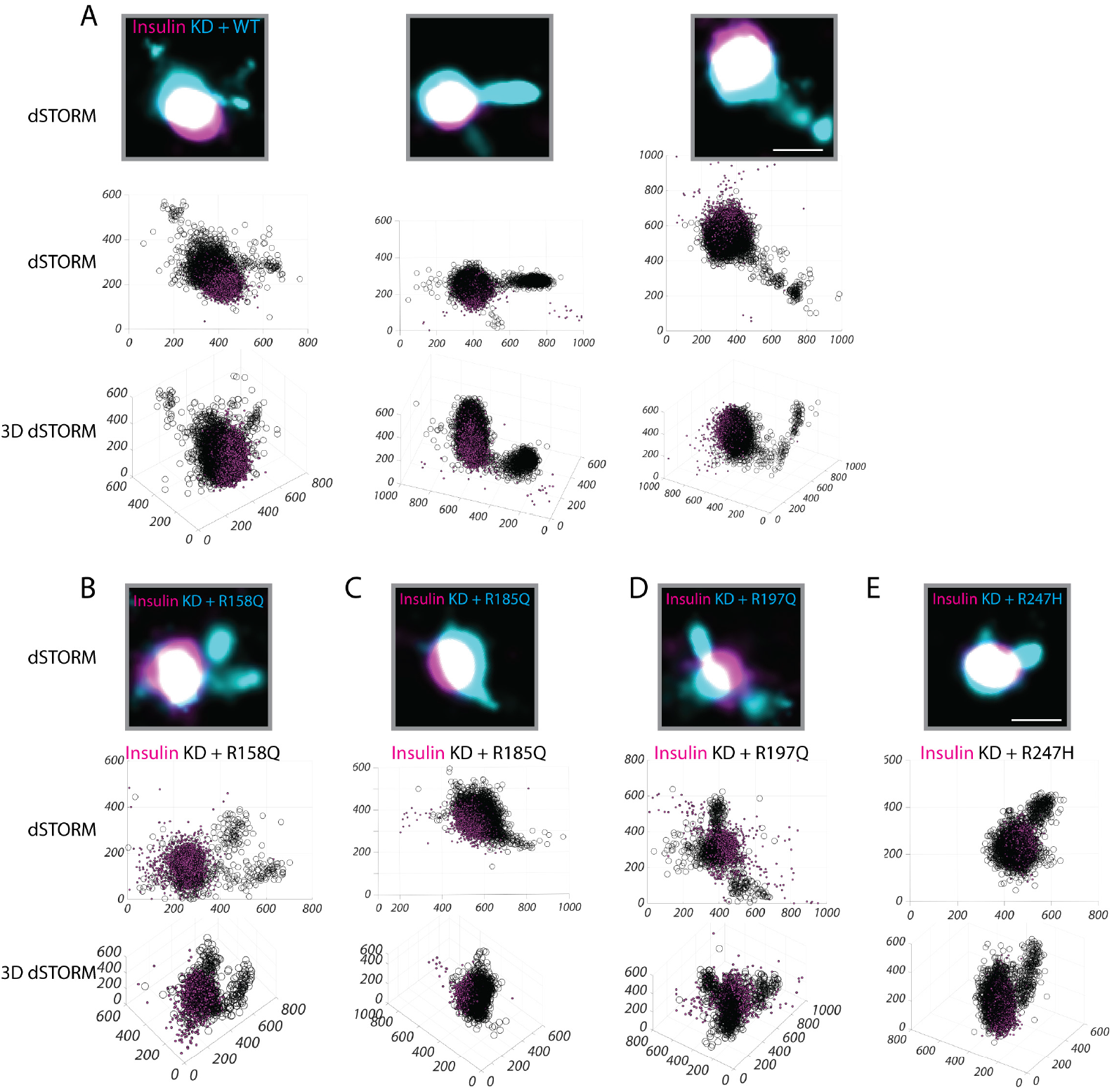
PICK1 WT and the coding variants localize to extensions from insulin granules. (A) INS-1E cells transduced with the lentiviral construct KD + WT. (C-F) INS-1E cells transduced with the PICK1 coding variants. (A-E) Upper row show representative dSTORM images of an insulin granule (magenta) colocalized with PICK1 (cyan) which is also extending from insulin granules. Scale bar = 250 nm. The middle row shows the xy structure of insulin (magenta) and PICK1 (black) and the lower row shows the 3D structure. Axis are in nm.

**Supplemental Figure 11 (Related to Figure 8).**
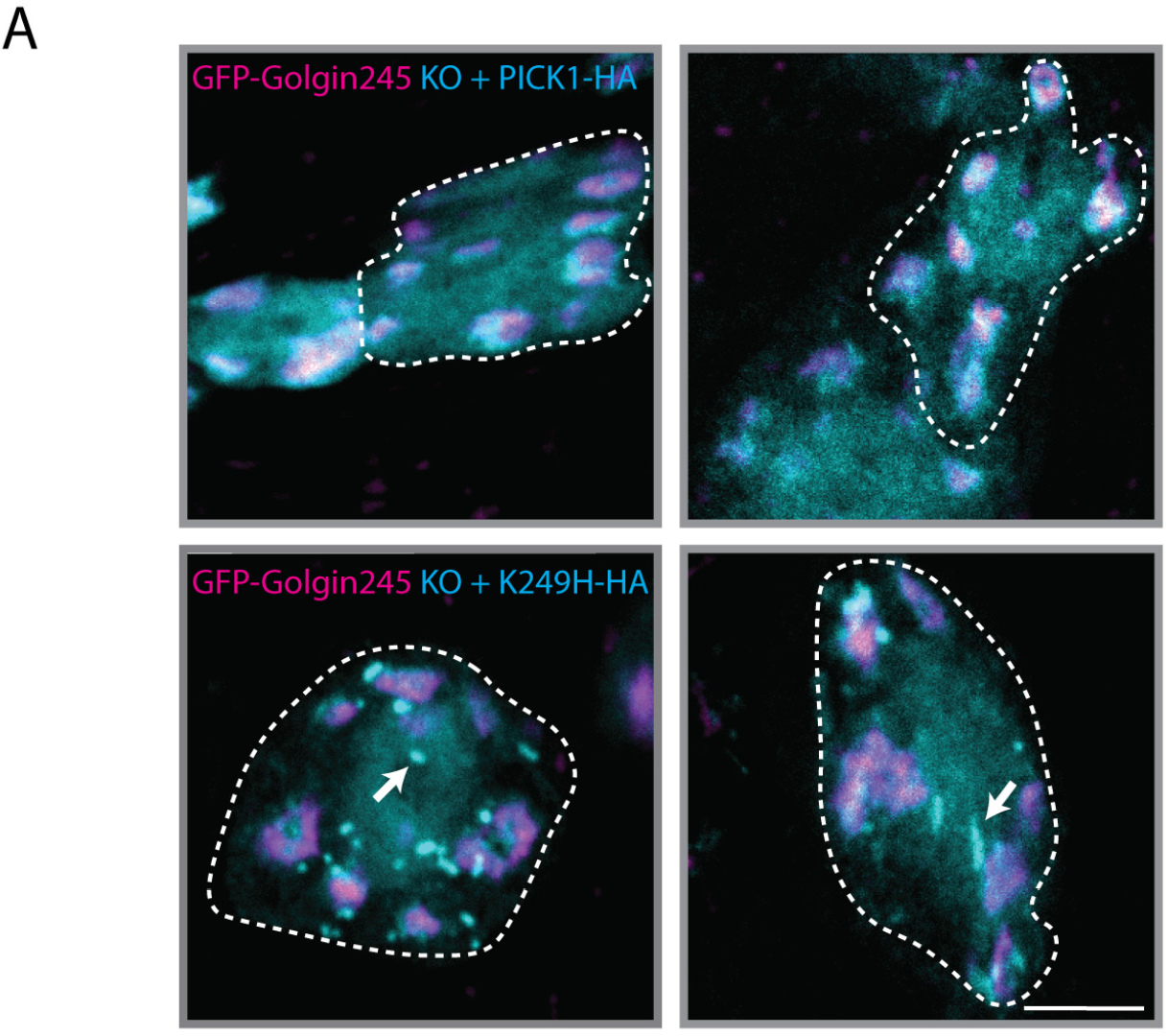
Perturbed subcellular distribution of *Drosophila* PICK1^K249H^ in peptidergic neurons. (A) Anti-HA immunolabeling of large peptidergic somata in the ventral nerve cord of pupal flies expressing the TGN marker GFP-Golgin245 (magenta) and either WT dPICK1-HA (*top*) or dPICK1K249H-HA (*bottom*) (cyan) in a dPICK1 null background. White dotted lines represent peptidergic neurons. dPICK1K249H-HA forms brightly labeled punctate clusters and tubules (arrows) in the cytosol that are not seen for WT dPICK1-HA. These structures remain tightly associated with the TGN. Scale bar = 5 µm.

## References

1. Zheng Y, Ley SH, and Hu FB. Global aetiology and epidemiology of type 2 diabetes mellitus and its complications. Nat Rev Endocrinol. 2018;14(2):88–98.

2. Ashcroft FM, and Rorsman P. Diabetes mellitus and the beta cell: the last ten years. Cell. 2012;148(6):1160–71.

3. Lin WJ, and Salton SR. The regulated secretory pathway and human disease: insights from gene variants and single nucleotide polymorphisms. Front Endocrinol (Lausanne). 2013;4:96.

4. Borgonovo B, Ouwendijk J, and Solimena M. Biogenesis of secretory granules.Curr Opin Cell Biol. 2006;18(4):365–70.

5. Kim T, Gondre-Lewis MC, Arnaoutova I, and Loh YP. Dense-core secretory granule biogenesis. Physiology (Bethesda). 2006;21:124–33.

6. Roder PV, Wong X, Hong W, and Han W. Molecular regulation of insulin granule biogenesis and exocytosis. Biochem J. 2016;473(18):2737–56.

7. Kuliawat R, Klumperman J, Ludwig T, and Arvan P. Differential sorting of lysosomal enzymes out of the regulated secretory pathway in pancreatic beta-cells. J Cell Biol. 1997;137(3):595–608.

8. Castle AM, Huang AY, and Castle JD. Passive sorting in maturing granules of AtT-20 cells: the entry and exit of salivary amylase and proline-rich protein. J Cell Biol. 1997;138(1):45–54.

9. Eaton BA, Haugwitz M, Lau D, and Moore HP. Biogenesis of regulated exocytotic carriers in neuroendocrine cells. J Neurosci. 2000;20(19):7334–44.

10. Arvan P, Kuliawat R, Prabakaran D, Zavacki AM, Elahi D, Wang S, et al. Protein discharge from immature secretory granules displays both regulated and constitutive characteristics. J Biol Chem. 1991;266(22):14171–4.

11. Orci L, Ravazzola M, Amherdt M, Perrelet A, Powell SK, Quinn DL, et al. The trans-most cisternae of the Golgi complex: a compartment for sorting of secretory and plasma membrane proteins. Cell. 1987;51(6):1039–51.

12. Tooze J, and Tooze SA. Clathrin-coated vesicular transport of secretory proteins during the formation of ACTH-containing secretory granules in AtT20 cells. J Cell Biol. 1986;103(3):839–50.

13. Molinete M, Irminger JC, Tooze SA, and Halban PA. Trafficking/sorting and granule biogenesis in the beta-cell. Semin Cell Dev Biol. 2000;11(4):243–51.

14. Klumperman J, Kuliawat R, Griffith JM, Geuze HJ, and Arvan P. Mannose 6-phosphate receptors are sorted from immature secretory granules via adaptor protein AP-1, clathrin, and syntaxin 6-positive vesicles. J Cell Biol. 1998;141(2):359–71.

15. Arvan P, and Castle D. Neuroendocrinology: regulated secretory cells go to the BAR for a bud. Nat Rev Endocrinol. 2013;9(8):443–4.

16. Peter BJ, Kent HM, Mills IG, Vallis Y, Butler PJ, Evans PR, et al. BAR domains as sensors of membrane curvature: the amphiphysin BAR structure. Science. 2004;303(5657):495–9.

17. Boucrot E, Pick A, Camdere G, Liska N, Evergren E, McMahon HT, et al. Membrane fission is promoted by insertion of amphipathic helices and is restricted by crescent BAR domains. Cell. 2012;149(1):124–36.

18. Bhatia VK, Madsen KL, Bolinger PY, Kunding A, Hedegard P, Gether U, et al. Amphipathic motifs in BAR domains are essential for membrane curvature sensing. The EMBO journal. 2009;28(21):3303–14.

19. Gehart H, Goginashvili A, Beck R, Morvan J, Erbs E, Formentini I, et al. The BAR domain protein Arfaptin-1 controls secretory granule biogenesis at the trans-Golgi network. Developmental cell. 2012;23(4):756–68.

20. Cao M, Mao Z, Kam C, Xiao N, Cao X, Shen C, et al. PICK1 and ICA69 control insulin granule trafficking and their deficiencies lead to impaired glucose tolerance. PLoS biology. 2013;11(4):e1001541.

21. Holst B, Madsen KL, Jansen AM, Jin C, Rickhag M, Lund VK, et al. PICK1 deficiency impairs secretory vesicle biogenesis and leads to growth retardation and decreased glucose tolerance. PLoS biology. 2013;11(4):e1001542.

22. Li J, Mao Z, Huang J, and Xia J. PICK1 is essential for insulin production and the maintenance of glucose homeostasis. Molecular biology of the cell. 2018;29(5):587–96.

23. Lohmueller KE, Sparso T, Li Q, Andersson E, Korneliussen T, Albrechtsen A, et al. Whole-exome sequencing of 2,000 Danish individuals and the role of rare coding variants in type 2 diabetes. Am J Hum Genet. 2013;93(6):1072–86.

24. Herlo R, Lund VK, Lycas MD, Jansen AM, Khelashvili G, Andersen RC, et al. An Amphipathic Helix Directs Cellular Membrane Curvature Sensing and Function of the BAR Domain Protein PICK1. Cell Rep. 2018;23(7):2056–69.

25. Spitzenberger F, Pietropaolo S, Verkade P, Habermann B, Lacas-Gervais S, Mziaut H, et al. Islet cell autoantigen of 69 kDa is an arfaptin-related protein associated with the Golgi complex of insulinoma INS-1 cells. J Biol Chem. 2003;278(28):26166–73.

26. Buffa L, Fuchs E, Pietropaolo M, Barr F, and Solimena M. ICA69 is a novel Rab2 effector regulating ER-Golgi trafficking in insulinoma cells. Eur J Cell Biol. 2008;87(4):197–209.

27. Jin W, Ge WP, Xu J, Cao M, Peng L, Yung W, et al. Lipid binding regulates synaptic targeting of PICK1, AMPA receptor trafficking, and synaptic plasticity. J Neurosci. 2006;26(9):2380–90.

28. Madsen KL, Eriksen J, Milan-Lobo L, Han DS, Niv MY, Ammendrup-Johnsen I, et al. Membrane localization is critical for activation of the PICK1 BAR domain. Traffic. 2008;9(8):1327–43.

29. Lu W, and Ziff EB. PICK1 interacts with ABP/GRIP to regulate AMPA receptor trafficking. Neuron. 2005;47(3):407–21.

30. Citri A, Bhattacharyya S, Ma C, Morishita W, Fang S, Rizo J, et al. Calcium binding to PICK1 is essential for the intracellular retention of AMPA receptors underlying long-term depression. The Journal of neuroscience : the official journal of the Society for Neuroscience. 2010;30(49):16437–52.

31. Li J, Barylko B, Eichorst JP, Mueller JD, Albanesi JP, and Chen Y. Association of Endophilin B1 with Cytoplasmic Vesicles. Biophys J. 2016;111(3):565–76.

32. Hatzakis NS, Bhatia VK, Larsen J, Madsen KL, Bolinger PY, Kunding AH, et al. How curved membranes recruit amphipathic helices and protein anchoring motifs. Nature chemical biology. 2009;5(11):835–41.

33. Zeno WF, Snead WT, Thatte AS, and Stachowiak JC. Structured and intrinsically disordered domains within Amphiphysin1 work together to sense and drive membrane curvature. Soft Matter. 2019;15(43):8706–17.

34. Snead WT, Zeno WF, Kago G, Perkins RW, Richter JB, Zhao C, et al. BAR scaffolds drive membrane fission by crowding disordered domains. J Cell Biol. 2019;218(2):664–82.

35. Karlsen ML, Thorsen TS, Johner N, Ammendrup-Johnsen I, Erlendsson S, Tian X, et al. Structure of Dimeric and Tetrameric Complexes of the BAR DomainProtein PICK1 Determined by Small-Angle X-Ray Scattering. Structure.2015;23(7):1258–70.

36. Haataja L, Snapp E, Wright J, Liu M, Hardy AB, Wheeler MB, et al. Proinsulin intermolecular interactions during secretory trafficking in pancreatic beta cells. J Biol Chem. 2013;288(3):1896–906.

37. Molinete M, Dupuis S, Brodsky FM, and Halban PA. Role of clathrin in the regulated secretory pathway of pancreatic beta-cells. J Cell Sci. 2001;114(Pt 16):3059–66.

38. Roux KJ, Kim DI, Raida M, and Burke B. A promiscuous biotin ligase fusion protein identifies proximal and interacting proteins in mammalian cells. J Cell Biol. 2012;196(6):801–10.

39. Kim DI, Jensen SC, Noble KA, Kc B, Roux KH, Motamedchaboki K, et al. An improved smaller biotin ligase for BioID proximity labeling. Molecular biology of the cell. 2016;27(8):1188–96.

40. Wu T, Shi Z, and Baumgart T. Mutations in BIN1 associated with centronuclear myopathy disrupt membrane remodeling by affecting protein density and oligomerization. PloS one. 2014;9(4):e93060.

41. Wise CA, Gillum JD, Seidman CE, Lindor NM, Veile R, Bashiardes S, et al. Mutations in CD2BP1 disrupt binding to PTP PEST and are responsible for PAPA syndrome, an autoinflammatory disorder. Hum Mol Genet. 2002;11(8):961–9.

42. McMahon HT, and Boucrot E. Membrane curvature at a glance. Journal of cell science. 2015;128(6):1065–70.

43. Simunovic M, Evergren E, Callan-Jones A, and Bassereau P. Curving Cells Inside and Out: Roles of BAR Domain Proteins in Membrane Shaping and Its Cellular Implications. Annu Rev Cell Dev Biol. 2019;35:111–29.

44. Madsen KL, and Herlo R. Recursive Alterations of the Relationship between Simple Membrane Geometry and Insertion of Amphiphilic Motifs. Membranes. 2017;7(1).

45. Wu T, and Baumgart T. BIN1 membrane curvature sensing and generation show autoinhibition regulated by downstream ligands and PI(4,5)P2. Biochemistry. 2014;53(46):7297–309.

46. Simunovic M, Mim C, Marlovits TC, Resch G, Unger VM, and Voth GA. Protein-mediated transformation of lipid vesicles into tubular networks. Biophys J. 2013;105(3):711–9.

47. Bonnemaison ML, Eipper BA, and Mains RE. Role of adaptor proteins in secretory granule biogenesis and maturation. Front Endocrinol (Lausanne). 2013;4:101.

48. Jansen AM, Nassel DR, Madsen KL, Jung AG, Gether U, and Kjaerulff O. PICK1 expression in the Drosophila central nervous system primarily occurs in the neuroendocrine system. J Comp Neurol. 2009;517(3):313–32.

49. Hewes RS, Park D, Gauthier SA, Schaefer AM, and Taghert PH. The bHLH protein Dimmed controls neuroendocrine cell differentiation in Drosophila. Development. 2003;130(9):1771–81.

50. Madsen KL, Beuming T, Niv MY, Chang CW, Dev KK, Weinstein H, et al. Molecular determinants for the complex binding specificity of the PDZ domain in PICK1. J Biol Chem. 2005;280(21):20539–48.

51. Erlendsson S, Rathje M, Heidarsson PO, Poulsen FM, Madsen KL, Teilum K, et al. Protein interacting with C-kinase 1 (PICK1) binding promiscuity relies on unconventional PSD-95/discs-large/ZO-1 homology (PDZ) binding modes for nonclass II PDZ ligands. The Journal of biological chemistry. 2014;289(36):25327–40.

52. Bolte S, and Cordelieres FP. A guided tour into subcellular colocalization analysis in light microscopy. J Microsc. u2006;224(Pt 3):213–32.

53. Ovesny M, Krizek P, Borkovec J, Svindrych Z, and Hagen GM. ThunderSTORM: a comprehensive ImageJ plug-in for PALM and STORM data analysis and super-resolution imaging. Bioinformatics. 2014;30(16):2389–90.

54. Malkusch S, and Heilemann M. Extracting quantitative information from single-molecule super-resolution imaging data with LAMA - LocAlization Microscopy Analyzer. Sci Rep. 2016;6:34486.

